# A cohesin acetylation complex that recognises nicked DNA

**DOI:** 10.64898/2026.11.05.764696

**Authors:** Céline Bouchoux, Frank Uhlmann

## Abstract

Eukaryotic DNA replication products are held together by cohesin to enable their faithful inheritance during cell division. The establishment of stable sister chromatid cohesion requires replication-coupled acetylation of two conserved lysines on cohesin’s Smc3 ATPase head. Here, we use AlphaFold 3 to predict the existence of a budding yeast ‘cohesin acetylation complex’, consisting of the replication protein PCNA, the cohesin subunit Pds5 and the Eco1 acetyltransferase. The predicted complex specifically contacts nicked DNA, a replication fork determinant that controls acetylation. Furthermore, cohesin engagement results in predicted conformational adaptations that bring acetyl-CoA in striking distance of the Smc3 target lysine. We perform biochemical and genetic experiments that confirm anticipated features of this reaction trajectory. Related cohesin acetylation complexes are predicted to form in other eukaryotes, including humans. Our study combines AI-assisted structure predictions and wet-bench experimentation to reveal the likely mechanism of replication-coupled cohesin acetylation during sister chromatid cohesion establishment.

## Introduction

Eukaryotic DNA replication is linked to inheritance of the two replication products by virtue of the chromosomal cohesin complex^1^. Cohesin, a member of the ring-shaped SMC complex family, topologically entraps the two newly synthesised sister chromatids. The resulting sister chromatid cohesion provides the mechanical counterforce to the mitotic spindle apparatus as it biorients chromosomes in preparation for chromosome segregation^2–4^. Replication-coupled sister chromatid cohesion establishment entails two essential aspects. Firstly, the two replication products must be co-entrapped inside individual cohesin rings. Recent work revealed that replisomes can pass through cohesin rings that have been loaded onto DNA before the onset of DNA replication, thus leaving replication products naturally co-entrapped^5^. This process, however, is successful in only a subset of replisome-cohesin encounters. Often, the cohesin ring transfers to only one of the two replication products, and it remains incompletely understood whether and how these cohesins capture the second sister chromatid^6–8^.

A second essential aspect of sister chromatid cohesion establishment is replication-coupled cohesin acetylation of two conserved lysine residues on its Smc3 ATPase head^9–11^. In budding yeast, acetylation is achieved by the Eco1 acetyltransferase, at first identified as a factor essential for establishment of sister chromatid cohesion, but not itself part of the cohesin complex^12–14^. Smc3 acetylation stops the otherwise dynamic cohesin loading and unloading cycles on and off DNA^15–17^, making this modification a key requirement for enduring sister chromatid cohesion. Cohesin is loaded onto DNA with help of the Scc2-Scc4 cohesin loader and is coupled to ATP hydrolysis. The two Smc3 ATPase head lysines contact both DNA and the cohesin loader in a central ATP and DNA-bound loading intermediate^18–20^. Acetylation precludes formation of this loading intermediate^21^, thus blocking cohesin loading onto DNA. Cohesin unloading from chromosomes, on the other hand, is promoted by ATP hydrolysis in conjunction with the Pds5-Wapl unloading complex. While the unloading mechanism remains incompletely understood, it likely involves a related ATP and DNA-bound transition state that involves the two Smc3 ATPase lysines.

Given that acetylation blocks both cohesin loading and unloading reactions^22^, this modification must be tightly regulated in space and time. Acetylation must wait until cohesin has successfully co-entrapped both sister chromatids behind the replication fork, but must then be swiftly completed before cohesin has a chance to unload following sister chromatid co-entrapment. Spatiotemporal control is at least in part achieved by the dependency of the cohesin acetylation reaction on transient DNA structures that arise during Okazaki fragment processing, DNA flaps and nicks^23^. These DNA structures define an acetylation window behind the replication fork, offering an explanation for how cohesin that has just co-entrapped two sister chromatids is stabilised.

How flaps and nicks are recognised and promote the cohesin acetylation reaction remains unknown. Biochemical reconstitution of replication-coupled cohesin acetylation revealed requirements additional to DNA structures. These include the replication sliding clamp PCNA, Scc2-Scc4 and Pds5, as well as the ability of cohesin to hydrolyse ATP^23^. Here, we use AlphaFold 3^24^ to probe how these components come together to jointly achieve DNA structure recognition and cohesin acetylation. We report the predicted structure of a cohesin acetylation complex consisting of PCNA, Pds5 and Eco1 that specifically engages nicked DNA. Addition of Smc3 and Scc1 causes conformational adaptations of the cohesin acetylation complex that bring acetyl-CoA in the Eco1 active site into catalytic distance of the Smc3 acetyl-acceptor lysine 113. We use biochemistry and genetics to explore these predictions, and we confirm key anticipated protein-protein and protein-DNA interactions that are instrumental for cohesin acetylation and sister chromatid cohesion establishment. Together, these approaches reveal the likely mechanism by which DNA replication is coupled to cohesin acetylation in order to stabilise newly established sister chromatid cohesion.

## Results

### A cohesin acetylation complex comprising PCNA, Pds5 and Eco1

Biochemical reconstitution of replication-coupled cohesin acetylation revealed the requirement for a DNA flap or nick structure that promotes the reaction^23^. Both structures share the discontinuity of one strand of the double helix as a defining feature. As the geometry of a nick is simpler than that of a flap, we here study how a nicked DNA substrate promotes cohesin acetylation. Using an electrophoretic mobility shift assay, we found that Eco1 readily and directly associates with DNA *in vitro*. However, by itself, Eco1’s affinity for a nicked DNA substrate was not measurably greater than that for plain double stranded DNA (Figure S1A). The ability to specifically engage with DNA in a structure-specific manner therefore likely involves components additional to Eco1.

To explore whether, and how, additional protein factors contribute to DNA structure recognition, we utilised AlphaFold 3^24^. We performed structure predictions of combinations of the protein components PCNA, Scc2-Scc4, Pds5, and Eco1 that together promote replication-coupled cohesin acetylation. As the DNA substrate, we used a 61 bp long double stranded DNA with a single nick at position 46. Eco1 contains a predicted zinc finger domain, so we included zinc, as well as the Eco1 substrate acetyl-CoA. Of all tested groupings, combining PCNA, Pds5, and Eco1 with nicked DNA and the co-factors resulted in the formation of a three subunit complex that engages the nicked DNA substrate (Figure 1A). We name this three-component protein assembly the ‘cohesin acetylation complex’.

**Figure 1.**
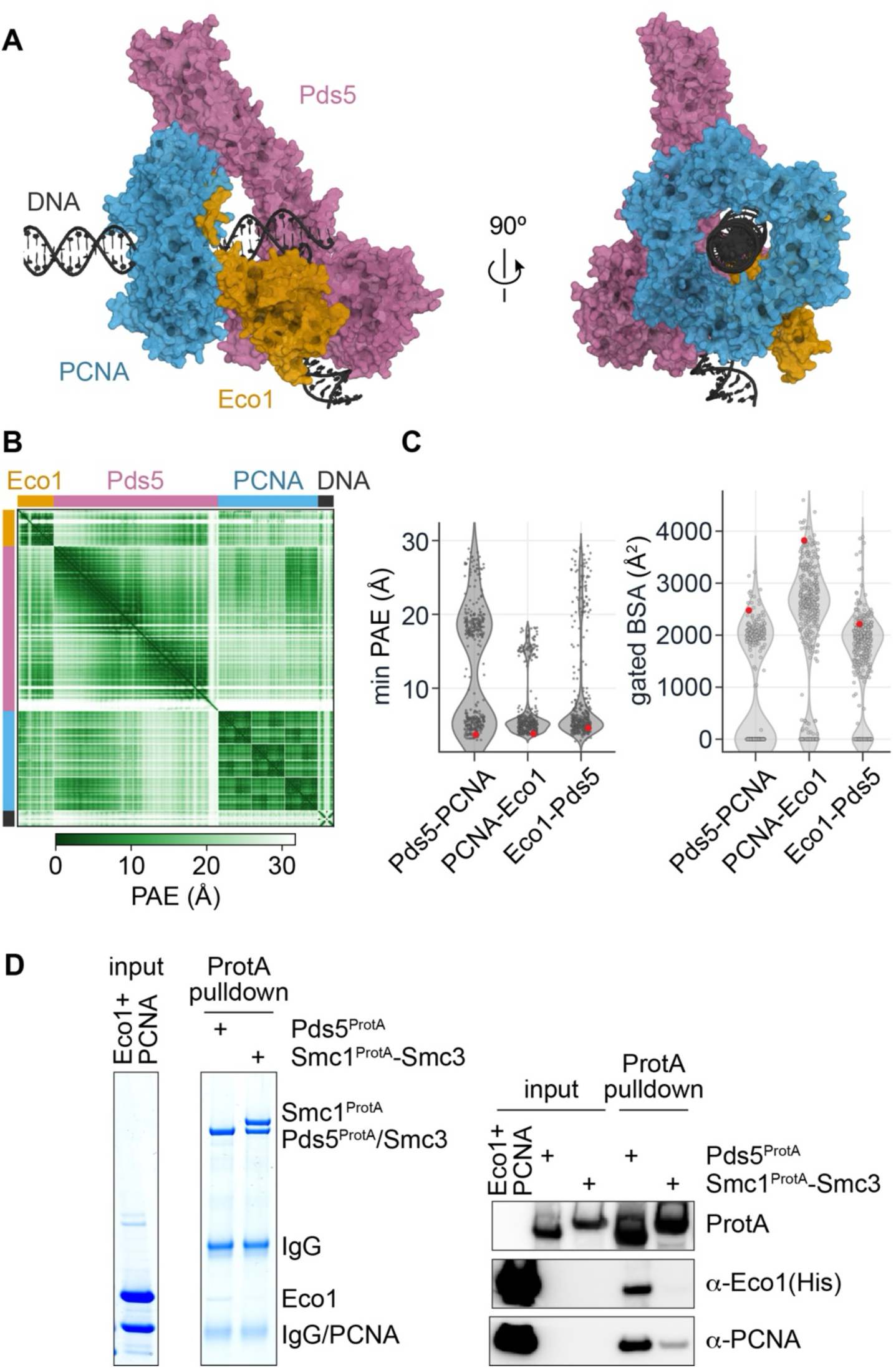
An AlphaFold 3 predicted cohesin acetylation complex comprising PCNA, Pds5 and Eco1 that engages nicked DNA. (A) AlphaFold 3 model (top ranked of those with a high Pds5-DNA buried surface area) of an *S. cerevisiae* PCNA-Pds5-Eco1 complex bound to nicked DNA (PCNA, blue; Pds5, pink; Eco1, gold; DNA, grey), shown in two views related by a 90° rotation. (B) Predicted aligned error (PAE) matrix of the cohesin acetylation complex model shown in (**A**). (C) Violin plots showing the spread of the minimum interchain PAE (min PAE) values, as well as the buried surface area (BSA) at the indicated protein interfaces amongst 375 generated models. The model shown in (A) is highlighted red. (D) Pull-down of purified PCNA and Eco1 by Pds5. Pds5 or Smc1-Smc3 as a negative control, fused to a protein A (ProtA) tag, were immobilised on IgG beads and incubated with recombinant Eco1 and PCNA. Input and pulldown fractions were resolved by SDS-PAGE, followed by Coomassie staining and Western blot analysis with the indicated antibodies.

With its predicted Local Distance Difference Test (pLDDT), AlphaFold 3 provides a per-residue confidence score of the prediction. The pLDDT score was high or very high throughout most of the expected structured parts of the three protein components, PCNA, Pds5 and Eco1^25–30^, but low or very low in parts of their expected unstructured regions (Figure S1B). In our structural models, we display residues assigned a pLDDT score of at least 50, thus avoiding the presentation of low confidence or disordered regions.

To analyse Alpha Fold 3’s confidence in predicted protein-protein interactions, we turned to the Predicted Aligned Error (PAE), the expected positional error in Ångströms (Å) between any pair of residues. The PAE matrix shows confident predictions of the protein domains that make up each of the components, indicated by PAE values of ∼ 5 Å. The three protein interfaces in turn showed interchain PAE values < 15 Å, again a sign of good confidence in the predicted subunit arrangement (Figure 1B).

To investigate the robustness of this AlphaFold 3 predicted cohesin acetylation complex, we generated 375 models (5 each from 75 seeds). Across these predictions, we plotted their minimum interchain PAE (min PAE) values as a measure for interaction confidence^31^, as well as the solvent-inaccessible buried surface area at the three subunit interfaces, considering only confidently assigned areas with an interchain PAE of < 15 Å (Figure 1C). This analysis revealed a min PAE value distribution that clustered around 5 Å at each interface, indicating confident prediction of the cohesin acetylation complex interactions. Buried surface areas were substantial and reached from ∼ 1,800 Å^2^ at the Pds5-PCNA interface to ∼ 2,500 Å^2^ between Eco1 and Pds5.

A direct protein interaction between PCNA and Pds5 was unanticipated, and in case of the PCNA-Pds5 interface we observed a subpopulation of models with distinctly greater min PAE, i.e. with a less confidently predicted Pds5-PCNA interface (Figure 1C). AlphaFold 3 assesses its models using a ranking score that combines measures of confidence in the overall structure, the predicted interfaces, as well as additional parameters, e.g. penalties for clashes. Those models with a greater min PAE at the Pds5-PCNA interface showed an overall lower ranking (Figure S1C). Conversely, the higher ranked AlphaFold 3 models present a closely aligned Pds5-PCNA interface, adding confidence in the prediction of this unanticipated interaction.

Given that not all models presented an equally well defined Pds5-PCNA interface, we explored whether AlphaFold can predict alternative conformations of the cohesin acetylation complex. To facilitate prediction of alternative states we used AlphaFold 3 with repulsive bias (AF3-ReD)^32^. Addition of a mathematical repulsive component into AlphaFold’s generative diffusion model can help predict alternate protein conformations. However, 375 models generated by AF3-ReD all closely resembled the cohesin acetylation complex predicted by AlphaFold 3 (Figure S1D). Taken together, our protein structure predictions suggest that PCNA, Pds5 and Eco1 engage in extensive interactions to form a three-component cohesin acetylation complex around a nicked DNA substrate.

We looked for experimental evidence of a cohesin acetylation complex in budding yeast. To do so we immobilised Pds5, fused to a Protein A affinity tag, onto IgG coupled magnetic beads. After incubation of these Pds5 beads with purified PCNA and Eco1, followed by extensive washes, both PCNA and Eco1 were retained, consistent with cohesin acetylation complex formation (Figure 1D). No, or much less, PCNA and Eco1 remained bound when we similarly immobilised Smc1-Smc3 dimers as a negative control. We also detected formation of a PCNA-Pds5-Eco1 complex in the presence of nicked DNA during glycerol gradient centrifugation in the presence of protein crosslinkers (GraFix^33^, Figure S1E). However, we were unable to capture the complex on a cryo-EM grid for structure determination.

### Experimental exploration of predicted subunit interfaces

#### Pds5-PCNA

AlphaFold 3 predicts two regions of contact between the two proteins, involving two characteristic Pds5 features that have no previously assigned functions (Figure 2A). Firstly, a peptide loop emanates from the HEAT repeats that has been referred to as the Pds5 ‘nose’^27,28^. This loop inserts into a hydrophobic groove on PCNA formed beneath its interdomain connecting loop, where otherwise PCNA-client protein interactions take place via a ‘PCNA interacting peptide’ PIP box motif (QxxΨxxΩΩ, where Ψ denotes an aliphatic and ΩΩ two aromatic residues)^34^. The Pds5 loop contains no sequence resembling a PIP box. Instead, the loop approaches the PCNA interdomain connector loop from a very different orientation and engages in a series of hydrophobic contacts that are largely distinct from those made by PIP box consensus residues (Figures 2A and S2A). An exception is the hydrophobic PCNA pocket that accommodates the PIP box Ψ residue, which is occupied by a Pds5 loop proline. The interchain PAE of the predicted interactions are < 5 Å, indicating good confidence in this interface (Figure S2B).

**Figure 2.**
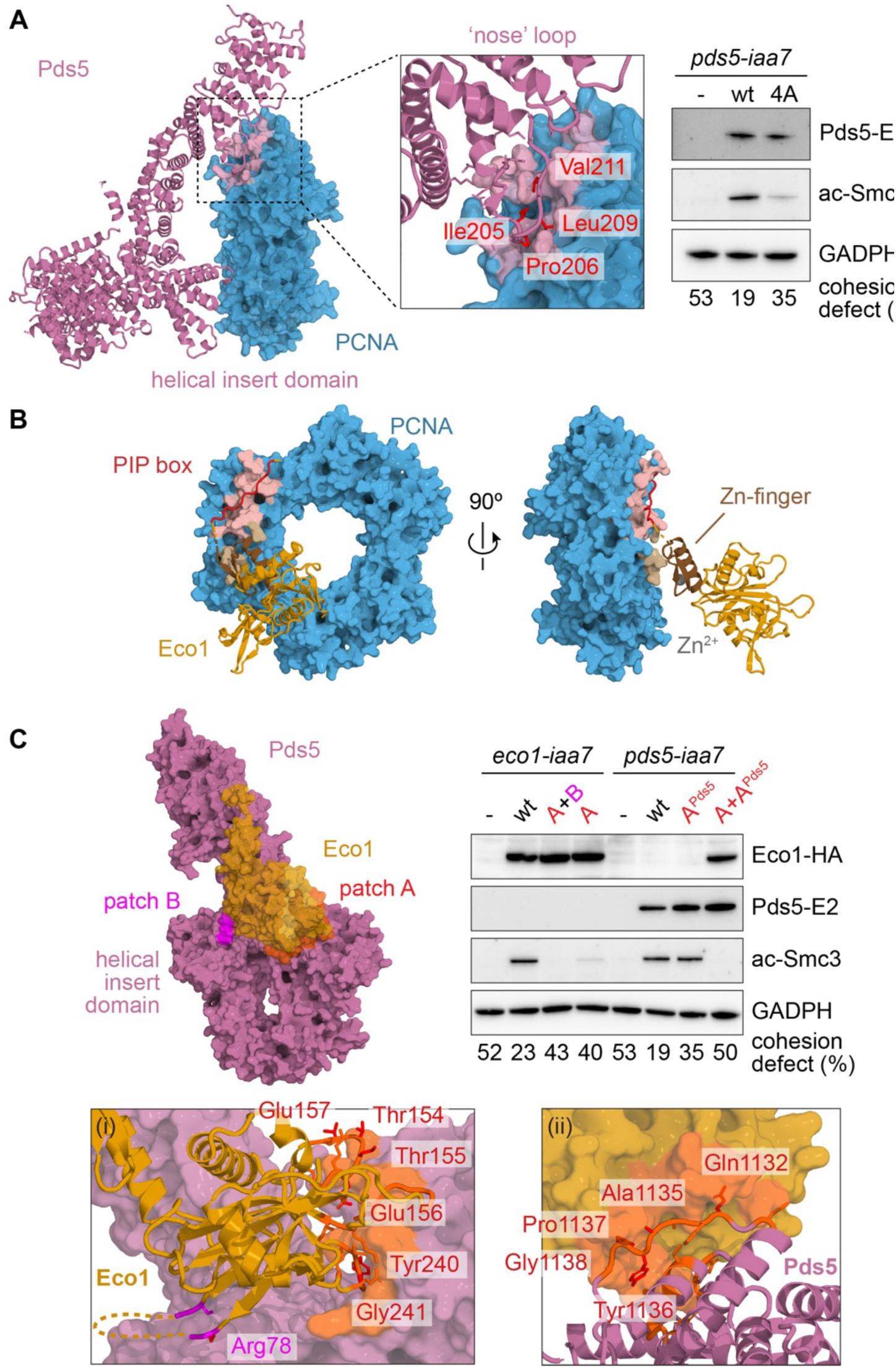
Protein interfaces of the cohesin acetylation complex. (A) The Pds5-PCNA interface. The Pds5 ‘nose’ loop and helical insert domains both contact PCNA. A close-up of the Pds5 ‘nose loop’ interaction is shown, mutated residues are highlighted in red. Yeast strains carrying an auxin inducible Pds5 degron (*pds5-iaa7*) were synchronised in G1 and released following auxin addition into a mitotic block, where the ability of ectopic wild type Pds5 and Pds5 4A (with the four indicated residues changed for alanines) to promote Smc3 acetylation and sister chromatid cohesion was assessed. (B) The PCNA-Eco1 interface, shown in two views related by a 90° rotation. (C) The Eco1-Pds5 interface, highlighting two interaction patches A (orange) and B (pink). Close-ups are shown of (i) patches A and B to highlight interacting residues on Eco1 and (ii) patch A to highlight interacting residues on Pds5. Mutated residues are highlighted in red. Yeast strains carrying auxin inducible Eco1 (*eco1-iaa7*) or Pds5 degrons were synchronised in G1 and released following auxin addition into a mitotic block, where the ability of ectopic wild type and mutant proteins to promote Smc3 acetylation and sister chromatid cohesion was assessed. The mutations were Eco1 ‘A’ (Y240A G241A), Eco1 ‘A+B’ (R78A T154A T155A E156A E157A Y240A G241A), Pds5 ‘A^Pds5^’ (Q1132A A1135D Y1136A P1137G G1138P).

The second Pds5-PCNA interaction takes place at Pds5’s distinctive helical insert domain that protrudes from the HEAT repeat scaffold^26,27^. Numerous hydrogen bond and electrostatic contacts are predicted between the helical insert domain and PCNA. These target the neighbouring PCNA protomer, adjacent to its interdomain connecting loop towards the lumen (Figures 2A and S2B). AlphaFold 3’s confidence in these interactions remained more tentative, with an interchain PAE above 15 Å. The interaction is nevertheless likely meaningful and is more confidently predicted in the context of additional components of the cohesin acetylation reaction (see below).

To test the relevance of a Pds5-PCNA interaction for sister chromatid cohesion establishment, we exchanged four hydrophobic residues in the Pds5 ‘nose’ loop to alanines (4A, Figure 2A). We then expressed Pds5 4A or wild type Pds5 in budding yeast cells from which we depleted endogenous Pds5 using an auxin inducible degron tag (*pds5-iaa7*)^35^. We synchronised cells in the G1 cell cycle phase using pheromone α-factor treatment, then added auxin to deplete endogenous Pds5 before releasing cells into a mitotic block imposed by the spindle poison nocodazole. We monitored cohesin acetylation using an acetyl-Smc3-specific antibody^36^, and the establishment of sister chromatid cohesion by observing the proximity of the GFP-marked *URA3* locus^37^. Pds5 depletion resulted in the absence of detectable cohesin acetylation and an observable sister chromatid cohesion defect in over half of cells. Smc3 acetylation was restored by expression of wild type Pds5, but to a substantially lower degree by Pds5 4A. Likewise, expression of wild type Pds5 reduced the occurrence of separate *URA3*-GFP signals to below 20%, while 35% of Pds5 4A expressing cells retained a sister chromatid cohesion defect. These observations suggest that a Pds5-PCNA interaction contributes to sister chromatid cohesin establishment in budding yeast.

#### PCNA-Eco1

PCNA was initially recognised as a high copy suppressor of the *eco1^ctf^*^7–203^ temperature sensitive mutation^13^. A direct physical PCNA-Eco1 interaction depends on a degenerate Eco1 PIP box motif (QxxLxxNN)^38^, where the two aromatic consensus residues are replaced with asparagines. Eco1 function in cohesin acetylation strictly depends on the consensus glutamine and leucine PIP-box residues^23^. Our AlphaFold 3 prediction illustrates how the degenerate Eco1 PIP box likely interacts with PCNA. The glutamine and leucine occupy the expected PIP box positions in the PCNA hydrophobic groove. In contrast, the two asparagines make contacts with the interdomain connecting loop, different from where consensus aromatic residues are known to bind (Figure S2A). These observations emphasise the versatility with which client proteins engage with PCNA.

In addition to the PIP box interactions, AlphaFold 3 predicts additional interactions between the Eco1 zinc finger and PCNA (Figure 2B). The zinc finger is also essential for replication-coupled cohesin acetylation^23,39^, but its molecular role remains unknown. To evaluate the relative contributions of the PIP box and zinc finger to the PCNA-Eco1 interaction we purified recombinant PCNA as well as wild type Eco1 and Eco1 variants with two alanine substitutions in the PIP box (Eco1 pip), or with the two zinc finger cysteines replaced by alanine (Eco1 zf). After combining PCNA and Eco1, we resolved the mixtures by size-exclusion chromatography. Both wild type Eco1 and Eco1 zf readily formed a complex with PCNA, but Eco1 pip did not (Figure S2C). This result suggests that the PIP box provides Eco1’s main affinity for PCNA. The zinc finger’s role likely lies in engaging cohesin as part of the cohesin acetylation reaction (see below).

#### Pds5-Eco1

Pds5 was identified in fission yeast as a high copy suppressor of the *eso1-H17* temperature sensitive mutation, and a yeast two-hybrid assay suggested that the two proteins interact^40^. Meanwhile, an AlphaFold-Multimer^41^ survey of sister chromatid cohesion factors predicted a direct budding yeast Pds5-Eco1 interaction^42^. Our AlphaFold 3 approach confirms and extends these findings (Figure 2C). In the predicted cohesin acetylation complex, Eco1 interacts with Pds5 at two places. An interaction patch A sits on the main Pds5 HEAT repeat body, as previously predicted^42^, while a second interaction patch B is situated on the helical insert domain.

To evaluate the importance of these Pds5-Eco1 contacts, we mutated residues on both sides of the predicted interface. On the Eco1 side, we replaced either Tyr240 and Gly241 of patch A to alanines (denoted ‘A’), or we mutated four additional patch A residues as well as the key arginine contact from patch B (‘A+B’, Figure 2C). The patch A only mutant showed a reduced ability to compensate for wild type Eco1 in cohesin acetylation and sister chromatid cohesion establishment, while the combined patch A and patch B mutant was almost completely defective in both activities. On the Pds5 side, we mutated complementary residues of the patch A interaction surface (denoted ‘A^Pds5^’). By themselves, they caused a moderate sister chromatid cohesion defect. Combining the ‘A^Pds5^’ mutations with the complementary Eco1 ‘A’ (Tyr240 and Gly241) changes resulted in a severe sister chromatid cohesion defect and complete loss of Smc3 acetylation. Together, these results suggest that a Pds5-Eco1 interaction makes an important contribution to cohesin acetylation and sister chromatid cohesion establishment.

### DNA nick recognition by the cohesin acetylation complex

To explore how the cohesin acetylation complex recognises its nicked DNA substrate, we performed AlphaFold 3 predictions of a PCNA-Pds5-Eco1 complex in the absence of DNA, or in the presence of undisrupted linear duplex DNA. A three component complex formed in each case. However, AlphaFold 3’s ranking scores for these complexes were substantially lower than for complexes assembled with nicked DNA (Figures 3A and S3A). Looking at 375 models with a nicked substrate, we find a bimodal distribution of DNA bending angles at the nick, with a majority (229) of models showing a strong bend of ∼ 74°, and the remainder displaying a slight curvature of ∼ 20° (Figure S3B). Bending went hand-in-hand with unwinding of 2 base pairs either side of the nick (Figure S3C). We then plot protein-DNA contacts made by the cohesin acetylation complex, defined as proximity of protein and DNA non-hydrogen atoms < 4.5 Å. Sequential contacts of PCNA, Eco1, and Pds5 are predicted along both linear and nicked DNA templates, but the frequency with which AlphaFold 3 predicts Pds5-DNA contacts increases in case of a nicked and bent DNA substrate (Figure 3B left). The buried surface area between Pds5 and DNA is low with linear DNA but increases to ∼ 2,300 Å^2^ in the case of nicked and bent DNA (Figure S3B). These findings suggest that the cohesin acetylation complex recognises a nicked DNA substrate by the sharp bend and local unwinding permitted by the discontinuity of the sugar-phosphate backbone. Furthermore, they suggest that the Pds5 subunit makes the main contribution to nick recognition.

**Figure 3.**
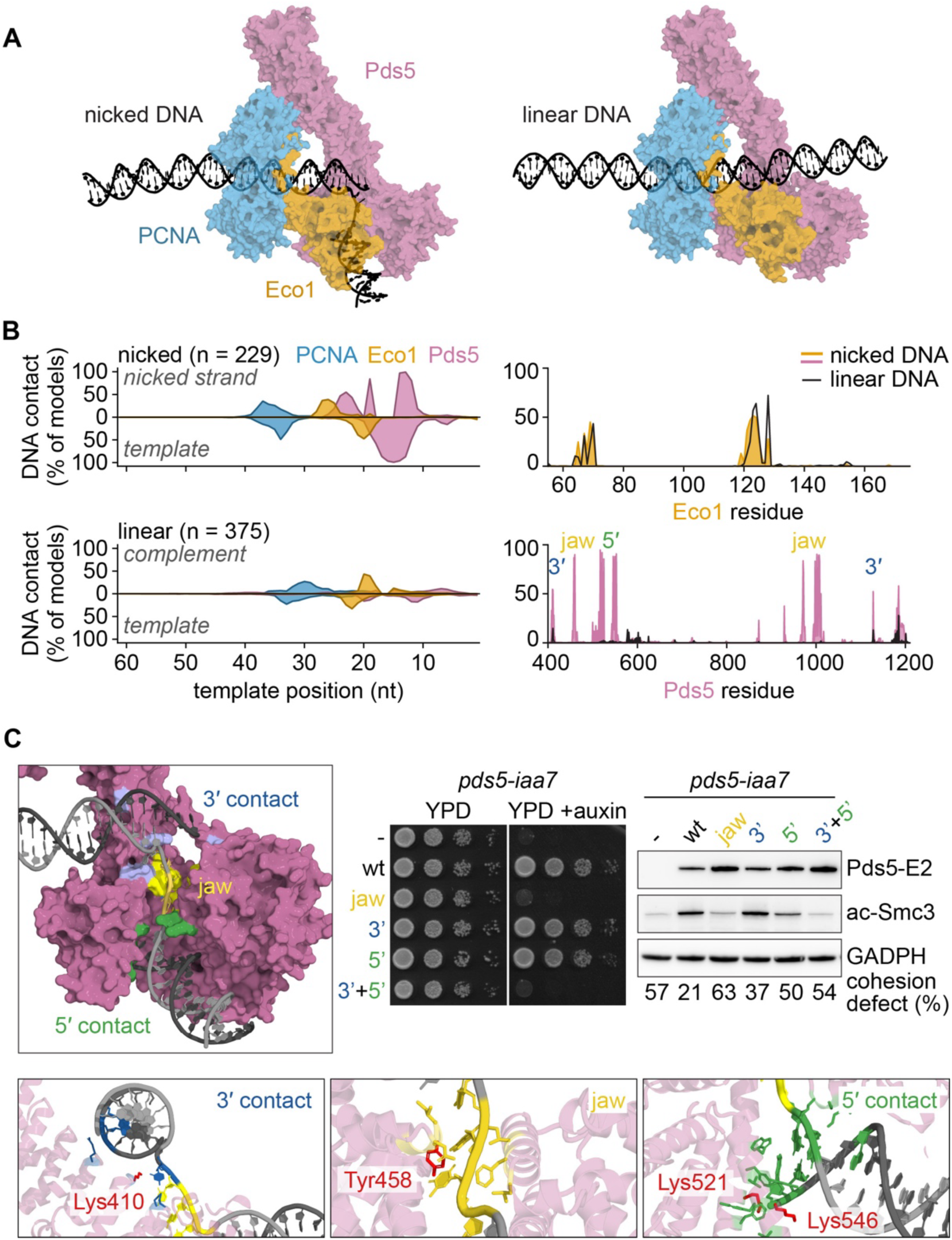
The cohesin acetylation complex engages nicked DNA. (A) Comparison of AlphaFold 3 predicted cohesin acetylation complexes formed with nicked or with linear DNA. The protein components are shown semi-transparent to highlight the DNA path. (B) Confidence-filtered DNA-contact footprints of PCNA, Eco1 and Pds5 along nicked (top left) and linear (bottom left) DNA. At each position, the fraction of models in which the protein makes a confident contact (protein-DNA heavy atom distance ≤ 4.5 Å, protein residue pLDDT > 50) is plotted along the template base-pairing register. Counterpart protein views of DNA-contacting residues in Eco1 (top right) and Pds5 (bottom right) are shown alongside. The Pds5 elements highlighted in panel (**C**) are indicated. (C) Close up of the Pds5 region predicted to engage the nick, as well as insets that give detail of how three aspects of the nick, the 3’ end, the unwound DNA (contacted by the jaw), and the 5’ end are engaged. Amino acids that were mutated are highlighted in red. Yeast strains carrying an auxin inducible Pds5 degron (*pds5-iaa7*) were synchronised in G1 and released following auxin addition into a mitotic block, where the ability of ectopic Pds5 with mutated jaw (Tyr457Ala Tyr458Ala), 5’ contact (Lys521Glu Lys546Glu), and 3’ contact (Lys410Glu His411Glu) to promote Smc3 acetylation and sister chromatid cohesion was assessed.

To experimentally probe nicked DNA recognition by Pds5, we mutated key residues predicted to engage with this DNA structure. Plotting DNA contacts along Pds5 identified three features predicted to engage the bent and locally unwound DNA (Figures 3B right and 3C). Firstly, the 3’ end of the nicked strand is held in place by charge interactions and a hydrophobic contact coming from the HEAT repeat surface, close to where the helical insert domain departs. We mutated Pds5 Lys410 and His411 that contact the DNA sugar-phosphate backbone at this place to glutamates. Second, the exposed single stranded region of the continuous DNA strand inserts into a jaw formed at the top of the Pds5 HEAT repeat crevice, where we mutated Tyr457 and Tyr458 that engage in stacking interactions with the exposed bases to alanines. Lastly, the 5’ end of the nicked strand is fixated by an array of hydrophobic and charge interactions emanating from both the HEAT repeats and the helical insert domain. Here we mutated Lys521 and Lys546 to glutamates. The jaw mutant and a combined 3’ and 5’ contact mutant displayed severe acetylation and sister chromatid cohesion defects after depletion of wild type Pds5, and were unable to support cell growth (Figure 3C). These results show that Pds5 residues that AlphaFold 3 predicts are involved in nicked DNA recognition are indeed important for cohesin acetylation.

In addition to Pds5, Eco1 also contributes DNA contacts in the predicted cohesin acetylation complex. While these contacts were equally observed with both linear and nicked DNAs (Figure 3B), we nevertheless tested whether they carry functional importance. Mutation of either Trp70 to alanine or Arg128 to glutamate substantially decreased the ability of Eco1 to perform its role in cohesin acetylation and sister chromatid cohesion establishment (Figure 4). These observations suggest that, while not involved in nick recognition, the ability of Eco1 to engage with DNA forms an important aspect of its function.

**Figure 4.**
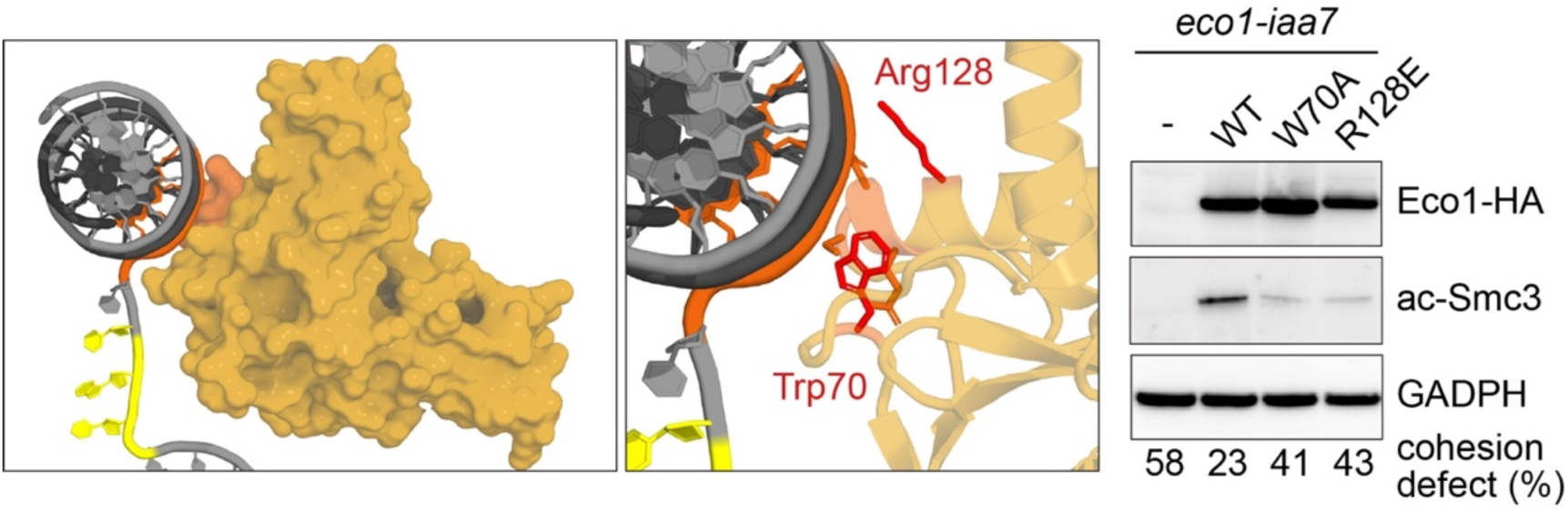
The Eco1 contact with DNA. Close up of Eco1 engaged with DNA, as well as inset that gives detail of the interaction. Amino acids that were mutated are highlighted in red. Yeast strains carrying an auxin inducible Eco1 degron (*eco1-iaa7*) were synchronised in G1 and released following auxin addition into a mitotic block, where the ability of ectopic Eco1 with Trp70Ala or Arg128Glu changes to promote Smc3 acetylation and sister chromatid cohesion was assessed.

While the cohesin acetylation complex forms an extensive protein-DNA interface in our predictions, the DNA in turn induces conformational adaptations that reshape protein-protein interactions. E.g. while a PCNA interaction with the Eco1 PIP box is observed in predicted cohesin acetylation complexes irrespective of the presence or absence of DNA, the zinc finger PCNA interaction is predicted only in the presence of DNA (Figure S4). The Pds5-Eco1 patch B interaction in turn is specific to the presence of nicked DNA. These observations suggest that nicked DNA is an integral component of cohesin acetylation complex assembly.

### The cohesin acetylation reaction

Next we included the Smc3 ATPase head and adjacent coiled coil in our AlphaFold 3 predictions, as well as the Scc1 N-terminus that connects the Smc3 coiled coil with Pds5^27,28,43^. The resulting predictions retained most features of the cohesin acetylation complex, with localised differences discussed below. We can see how Scc1 departs from Smc3 and inserts into the Pds5 HEAT repeat crevice as expected from the experimental structures. Notably, the Smc3 ATPase head is positioned right next to Eco1 (Figure 5A). While the same nicked DNA remains the template, its striking bend is no longer visible, and the path now resembles that of a linear DNA bound by the cohesin acetylation complex. The overall confidence in this predicted arrangement is very high, as seen from Alphafold 3 ranking scores for 375 models that surpass those of the three component cohesin acetylation complex (Figure S3A), as well as low interchain PAE values (Figure S5A).

**Figure 5.**
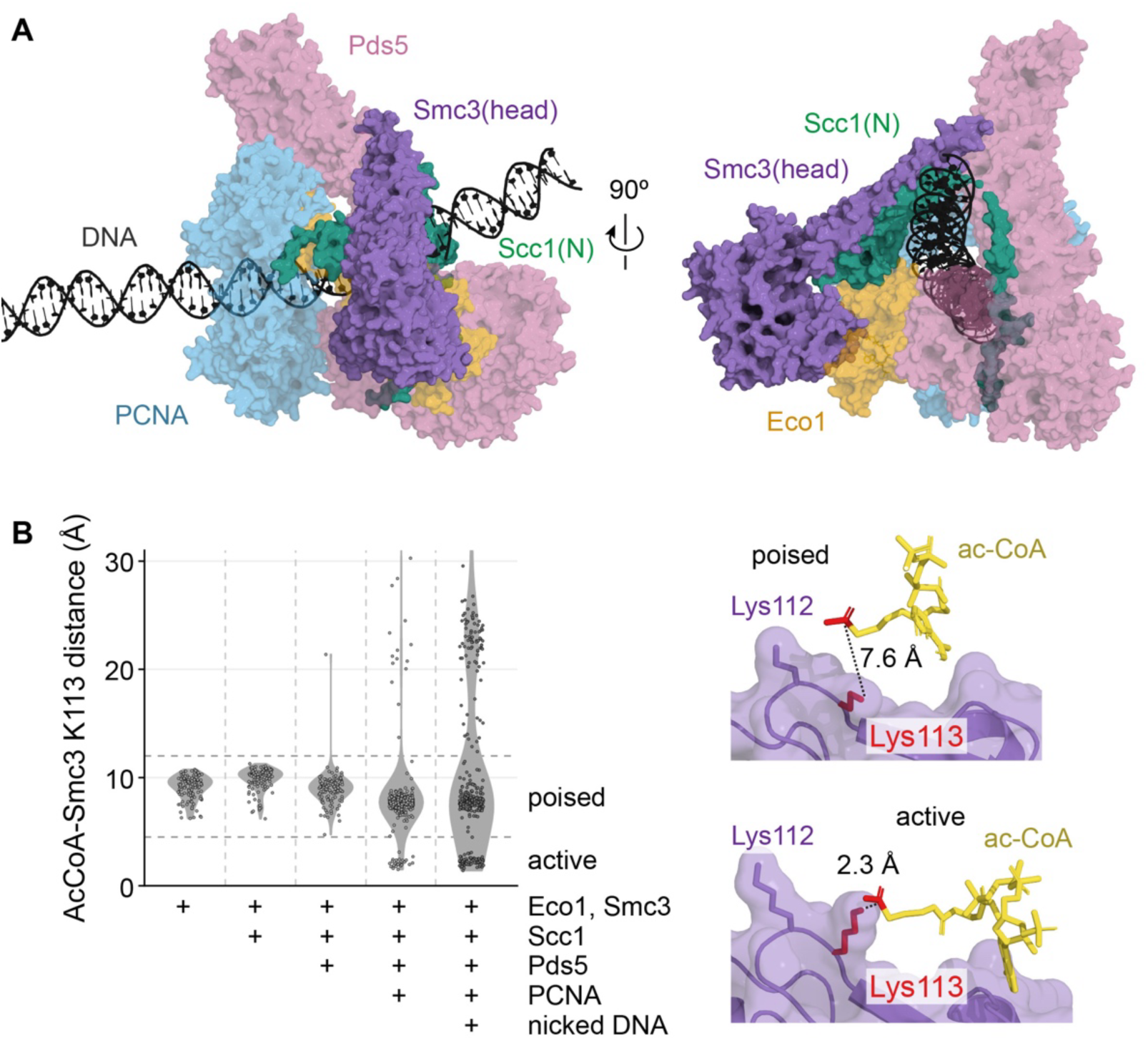
Smc3 Lys113 acetylation by the cohesin acetylation complex. (A) AlphaFold 3 model (top ranked of 375) of the *S. cerevisiae* PCNA-Pds5-Eco1 cohesin acetylation complex engaged with an Smc3 head domain plus 35 amino acids of adjacent coiled coil, Smc3(head), and a 151 amino acids Scc1 N-terminal portion, Scc1(N), bound to nicked DNA (PCNA, blue; Pds5, pink; Eco1, gold; Smc3, purple; Scc1, green; DNA, grey). Shown are two views related by a 90° rotation. (B) Distance between the carbonyl carbon of acetyl-CoA and the ζ-nitrogen in the Smc3 acceptor Lys113 in 125 AlphaFold 3 models of the indicated components (375 models each of the full complexes without and with DNA). Two distance regimes are indicated, ‘poised’ where Lys113 remains at a distance 4.5 - 12 Å, and ‘active’ where Lys113 has come within 4.5 Å of acetyl-CoA, close enough for acetyl transfer. Two example models of the poised and active state are shown.

AlphaFold multimer previously predicted that Eco1 engages the Smc3 ATPase head in an orientation that positions the Smc3 acetyl acceptor Lys113 side chain close to its active site^42^. This relative orientation is again seen when Eco1 is part of the cohesin acetylation complex. Moreover, as part of the cohesin acetylation complex the Eco1-Smc3 interaction experiences conformational adaptations that are likely critical for catalysis. When we measure the distance between the carbonyl carbon of acetyl-CoA and the ζ-nitrogen in the acceptor Lys113, the atom pair that will engage in chemical bond formation, we find three populations amongst the predictions. A majority population shows distances in the 10 Å range, consistent with predicted two-protein Eco1-Smc3 encounters (Figure 5B)^42^. At this distance acetyl-CoA is close to its target, but not close enough for acetyl transfer, and we refer to this configuration as the ‘poised’ configuration of the acetyltransferase. A small population of models shows far greater acetyl-CoA to Lys113 distances (> 20 Å), in which DNA has displaced Eco1 from Smc3. More notably, a subset of models bring acetyl-CoA within ∼2.5 Å of Lys113, a distance that enables acetyl transfer^44^. We call this arrangement the ‘active’ cohesin acetylation complex. This ‘active’ complex is observed only when all complex components, including PCNA, come together in the predictions, emphasising the likely importance of acetylation complex formation for catalysis.

To explore how the cohesin acetylation complex reaches its active state, we inspected Eco1’s predicted protein surroundings. Comparing the position of the Eco1 zinc finger, we notice a striking change between models in the poised and in the active states. In the poised configuration, the zinc finger is close to PCNA, not far from where it was located without Smc3 and Scc1 (Figure 6A). In contrast, the zinc finger is strikingly repositioned in the active state, now engaging Scc1 close to where Scc1 binds Smc3, as well as two surface loops on Smc3 (Figures S5B and S5C). Mutation or deletion of one of these Smc3 surface loops has recently been found to be detrimental for sister chromatid cohesion^45^, and a role in Eco1 zinc finger engagement could explain this observation.

**Figure 6.**
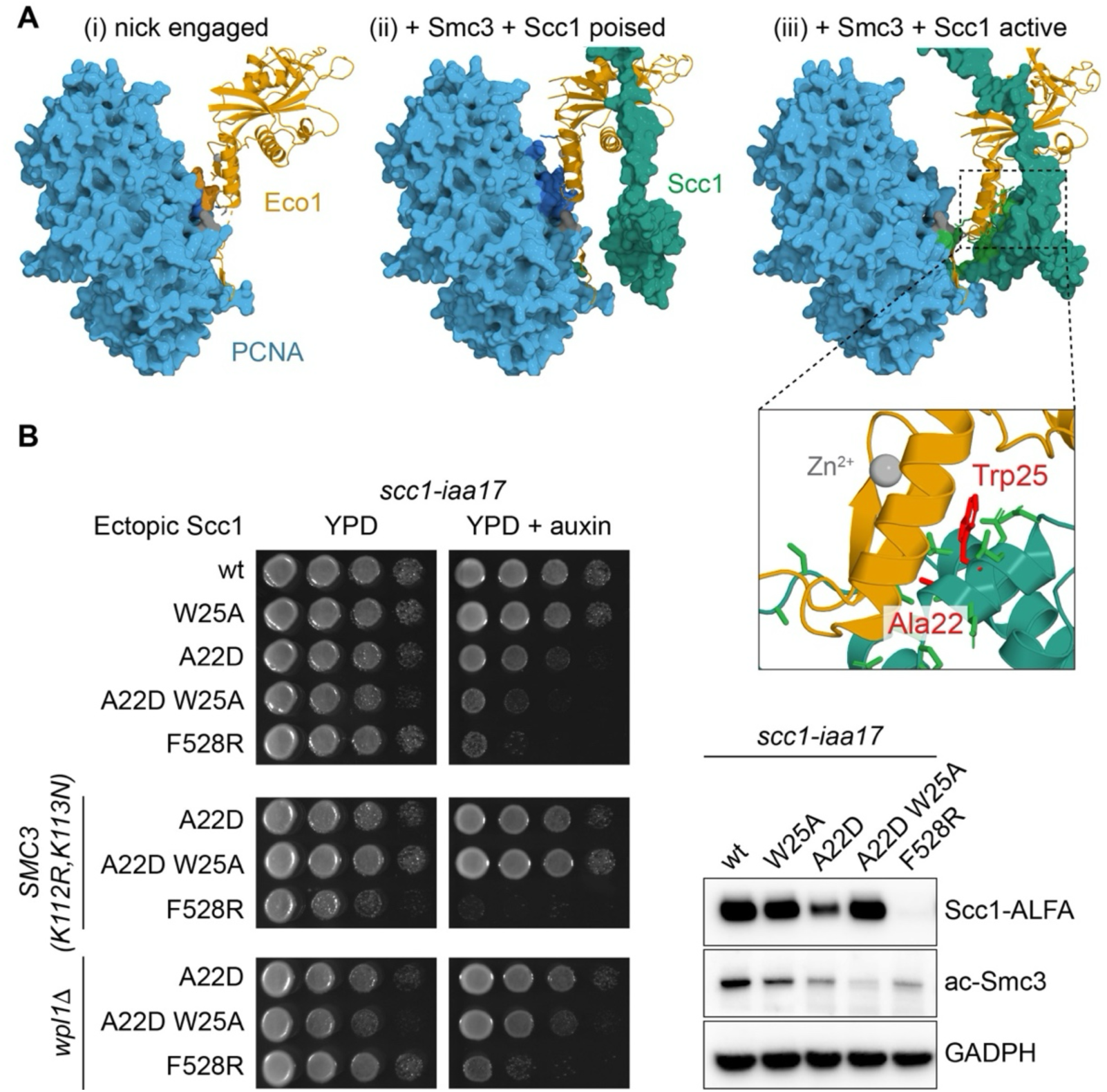
An Eco1 zinc finger-Scc1 interaction defines the acetylation active state. (A) Comparison of the predicted arrangement of Eco1 and its zinc finger, relative to PCNA and Scc1 in three states of the cohesin acetylation complex, (i) engaged with nicked DNA, (ii) in its poised state following addition of Smc3(head) and Scc1(N), (iii) in its active state. A close-up of the zinc finger-Scc1 interaction is shown, mutated residues are highlighted in red. (B) Yeast strains carrying an auxin inducible Scc1 degron (*scc1-iaa17*) carrying the indicated ectopic copies of Scc1 were spotted in 10-fold serial dilutions on YPD plates without or with auxin to deplete endogenous Scc1 to assess cell viability. Cells of all genotypes were grown on the same plate, separated solely to ease comparison. Alongside, cells were synchronised in G1 and released following auxin addition into a mitotic block, where the ability of ectopic Scc1 carrying the indicated mutations to promote Smc3 acetylation was assessed.

To test whether the predicted zinc finger-Scc1 contact participates in Smc3 acetylation we mutated two key residues of the interaction, Scc1 Ala22 to aspartate and Trp25 to alanine. Following depletion of endogenous Scc1, expression of both variants compromised cell viability and reduced Smc3 acetylation, defects that were exacerbated when the two changes were combined (Figure 6B). This result suggests that residues in Scc1 predicted to engage the Eco1 zinc finger to reach an acetyl transfer competent state are critical for *in vivo* cohesin acetylation.

Given the importance of the acetylation active state prediction, we tested whether the Scc1 Ala22 and Trp25 mutations specifically affect Smc3 acetylation, rather than in another way compromise cohesin function. If these mutations only disrupt Smc3 acetylation, they should become inconsequential in strain backgrounds where cohesin acetylation is dispensable. We tested this prediction in two ways. Firstly, we used an *SMC3(K112R,K113N)* strain background carrying an Smc3 variant in an unmodifiable but cohesion-permissive state. In this strain background the Scc1 Ala22 and Trp25 mutations no longer affected cell viability (Figure 6B). As a control, the growth defect caused by a Scc1F528R change, affecting the Scc1-Smc1 interaction^46^, remained unchanged in the *SMC3(K112R,K113N)* background. Secondly, if Smc3 acetylation is the reason for compromised cell growth, removing the cohesin unloader Wapl should ameliorate the growth defect^9^, which was indeed the case.

In further support of an Scc1 role in Smc3 acetylation, we found that Eco1 did not detectably interact with an Smc1-Smc3 heterodimer complex, but efficiently interacted with an Smc1-Smc3-Scc1 trimer complex (Figure S6A). *In vitro* Smc3 acetylation in a replication-coupled assay also strictly required Scc1 (Figure S6B), an observation consistent with results obtained with human proteins^47^. Together, these results suggest that AlphaFold 3 identified an unanticipated Eco1 zinc finger interaction with Scc1 that is instrumental in positioning Eco1 for acetyl transfer to the Smc3 Lys113 acceptor lysine.

### Additional considerations

Not all models of the cohesin acetylation complex in its active subunit configuration, i.e. with the Eco1 zinc finger engaging Scc1 and Smc3, showed the expected short acetyl-CoA to Smc3 Lys113 distance. In around half of the models Smc3 Lys113 instead pointed towards its neighbouring Smc3 acidic residue Asp115 (Figure S6C). A substrate-aided catalysis mechanism for Eco1 catalytic activity has been proposed in which a close-by acidic residue serves as the general base that helps extract a proton from Lys113 to aid its nucleophilic attack of the acetyl-CoA carbonyl^29^. It appears that Alphafold 3 has placed Lys113 in the active cohesin acetylation complex with approximately equal probability next to either a candidate general base or next to the acetyl-CoA carbonyl. During the actual catalysis cycle the three components likely adopt an intermediate configuration, achieving the required general base-assisted near attack distance for acetyl transfer of ∼3.5 Å^44^.

We earlier noted a relatively high PAE score, i.e. low confidence, of the Pds5 helical insert domain interaction with PCNA in the three component cohesin acetylation complex. If we inspect this interaction again in the active complex engaged with Smc3 and Scc1, we find that the PAE score at this interface has now substantially improved (Figure S2B). This observation suggests that the Pds5 helical insert domain adds to the conformational optimisation of the cohesin acetylation complex in preparation for acetyl transfer.

Inspecting the Scc1 path along the Pds5 crevice, we see that it overlaps with the predicted contact that the 5’ DNA end makes during nick recognition, close to the Pds5 helical insert domain (Figure S6D). If the prediction of how Pds5 engages nicked DNA is correct, this observation implies that the DNA displaces an Scc1 contact at this site during DNA substrate recognition, and that a handover back to Scc1 might happen as part of the cohesin acetylation reaction sequence. Pds5 is known to interact with cohesin in a dynamic fashion^16^, which might facilitate such exchanges.

### Conservation of the cohesin acetylation complex

Cohesin acetylation during sister chromatid cohesion establishment is a conserved feature amongst eukaryotes studied, including the fission yeast *S. pombe*^48^ and human^11,49,50^. We therefore explored whether cohesin acetylation complexes similar to the one in budding yeast are predicted to form across evolution. The fission yeast cohesin acetyltransferase Eso1 is a composite protein that consists of an Eco1-like acetyltransferase domain that makes up the C-terminal third, while the N-terminal two thirds are homologous to translesion DNA polymerase η^51^. Fission yeast cells appear to harbour two forms of Eso1, the full-length protein, as well as a proteolytically processed form with its two parts separated^52^. We first performed AlphaFold 3 predictions of the separated Eso1 acetyltransferase domain together with *S. pombe* PCNA and Pds5 in the presence of nicked DNA. This approach resulted in the prediction of a cohesin acetylation complex very similar to that observed with budding yeast (Figures 7A and S7A). The nicked DNA adopted a sharp bend and was partly unwound, suggesting that the fission yeast cohesin acetylation complex recognises its DNA substrate similarly to its budding yeast counterpart. A notable addition is a predicted ‘YSR’ motif interaction between an unstructured N-terminal extension of the Eso1 acetyltransferase domain and a Pds5 pocket to which also the Pds5 interactors Wapl and Sororin are known to bind. The contacts predicted to be made by the Eso1 ‘YSR’ motif peptide closely resemble those seen in an experimental structure of human WAPL bound to Pds5B^26^ (Figure 7B).

**Figure 7.**
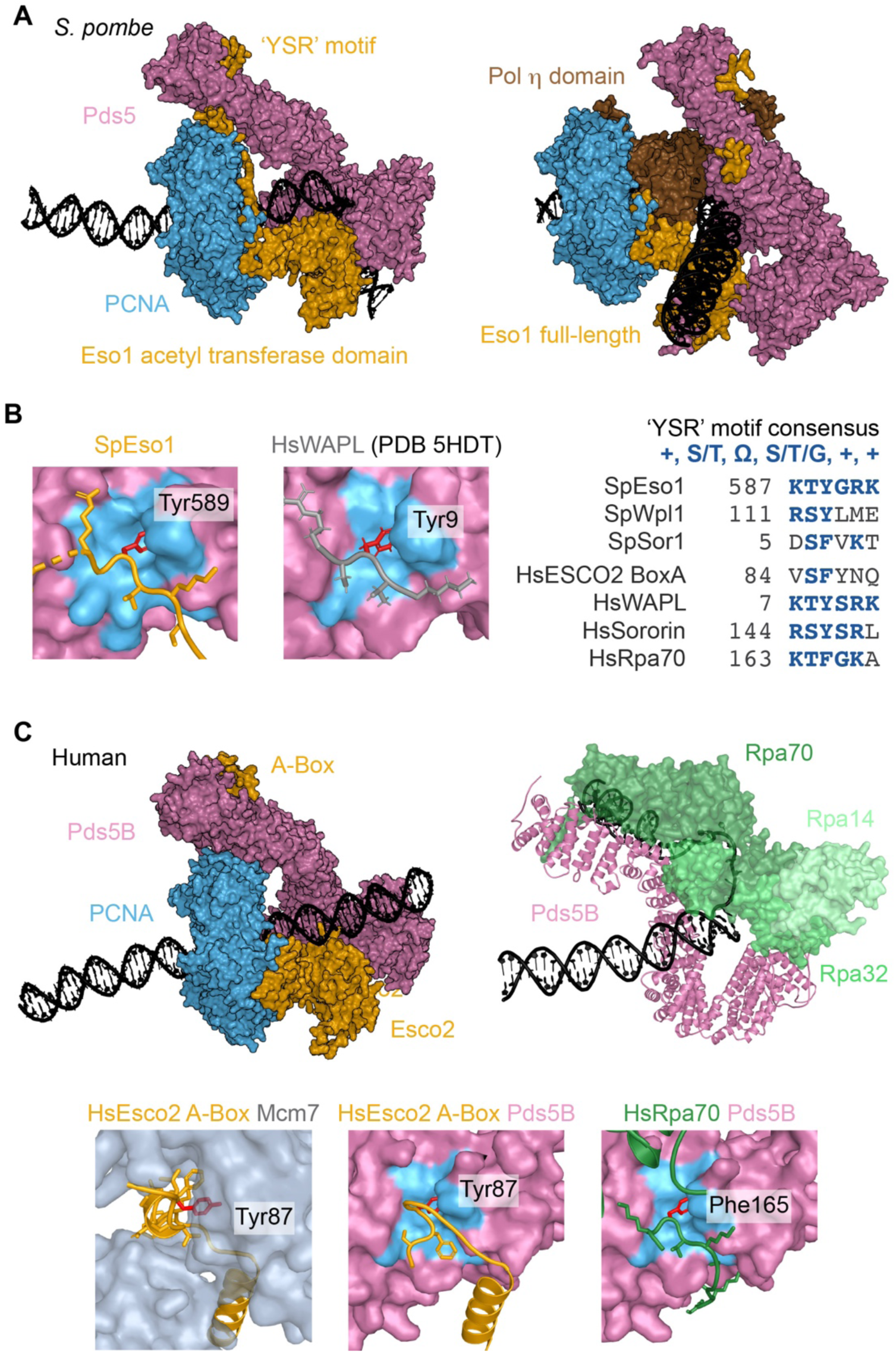
Predicted conservation of the cohesin acetylation complex. (A) AlphaFold 3 models (top ranked of those displaying bent DNA) of fission yeast *S. pombe* cohesin acetylation complexes formed of PCNA, Pds5, and either the Eso1 acetyltransferase domain or full-length Eso1 including its DNA polymerase η domain (PCNA, blue; Pds5, pink; Eso1 acetyltransferase domain, gold; Eso1 pol η domain, brown; nicked DNA, grey). (B) Close-up views of the Eso1 ‘YSR’ motif and its predicted binding to Pds5, next to the experimental structure of the human WAPL ‘YSR’ motif bound to Pds5B^26^. An alignment of ‘YSR’ motifs in the indicated proteins is included. (C) AlphaFold 3 model (top ranked of those showing Esco2 A-box engagement with Pds5B) of a human cohesin acetylation complex formed of PCNA, Pds5B, and Esco2 (PCNA, blue; Pds5B, pink; Esco2, gold; nicked DNA, grey). A model that includes the human trimeric RPA complex (green) is shown alongside formed around a 30 nt gapped DNA. Only Pds5B, RPA and DNA are shown for clarity. Close-up views of the predicted binding of the Esco2 A-box to MCM7 or Pds5B are shown, as well as of the RPA70 ‘YSR’ motif to Pds5B.

When using full-length Eso1, AlphaFold 3 predicts a very different arrangement. The DNA polymerase η domain replaced Pds5 to make extensive contacts with two PCNA protomers, while the acetyltransferase domain retained its own PCNA association. Thus, PCNA becomes entirely occupied by Eso1 (Figure 7A). In this assembly, DNA polymerase η is predicted to engage the nicked DNA by forming a bend, as it does during elongation of a primed DNA template^53^. This observation suggests that Eso1 can form part of two types of cohesin acetylation complexes, that each recognise nicked DNA by its ability to bend, yet in distinct ways. We note that the budding yeast DNA polymerase η contributes to DNA damage-induced sister chromatid cohesion establishment^54^. It could therefore be that DNA polymerase η forms part of cohesin acetylation complexes that acetylate cohesin as part of the DNA damage response^55^.

Finally we explored whether a cohesin acetylation complex also forms in humans. Human genomes encode two Pds5 isoforms that act largely redundantly^56^ and we display AlphaFold 3 predictions using Pds5B because of the available experimental structural comparison data^26^. Of the two human Eco1 orthologs, Esco1 and Esco2, we chose Esco2 that is responsible for replication-coupled cohesin acetylation and carries a conserved, essential PIP box^57,58^. Together with human PCNA, AlphaFold 3 predicts that the three components assemble into a cohesin acetylation complex with a conserved overall subunit arrangement (Figures 7C and S7A). Pds5B interacts with PCNA using both the ‘nose’ loop as well as the helical insert domain, while Esco2 uses residues conserved in both yeasts to engage with Pds5B (Figure S7B). Human Esco2 is also predicted to occupy the Pds5 ‘YSR’ motif receptor (Figure 7C). However, the Esco2 sequence diverges from a canonical ‘YSR’ motif. The divergent ‘YSR’ motif is part of the vertebrate-specific, conserved Esco2 ‘A-Box’^59^, which also mediates an Esco2 interaction with the MCM replicative helicase^60,61^. AlphaFold 3 predicts that Esco2 residues that interact with MCM7 overlap with those that bind Pds5. The MCM interaction is known to stabilise the Esco2 protein^60^, and it is possible that Esco2 is handed over from the MCM helicase to Pds5 in order to fulfil its essential role in sister chromatid cohesion establishment.

Even though the subunit arrangement of the human cohesin acetylation complex is predicted to resemble the yeasts, the way in which it engages with nicked DNA might differ. None of 375 models showed the nicked DNA in its characteristic bent configuration. Of the budding yeast Pds5 features that engage the nick, we find that only residues that occupy the 3’ end of the nicked strand and the beginning of the unwound strand (the top of the jaw) are conserved in human Pds5B (Figure S7C). Residues of the other half of the jaw, or that recognise the 5’ end, are not conserved. Instead, AlphaFold 3 predicted that the human single stranded DNA binding protein RPA interacts with Pds5, again including a ‘YSR’ motif interaction made by its RPA70 large subunit (Figure 7C). This observation opens the possibility that human cohesin acetylation is promoted by single stranded gaps that are decorated by RPA during Okazaki fragment synthesis.

## Discussion

In this study, we used AlphaFold 3 to predict the existence of a cohesin acetylation complex consisting of PCNA, the cohesin subunit Pds5, and the Eco1 acetyltransferase that assembles around nicked DNA, serving as the replication fork determinant for cohesin acetylation. AlphaFold 3 furthermore predicts how engagement of the cohesin acetylation complex with Smc3 and Scc1 brings conformational adaptations that guide acetyl-CoA in the Eco1 active site into striking distance from the Smc3 acetyl acceptor lysine. We experimentally explore these predictions and find that anticipated protein-protein and protein-DNA contacts play critical roles in the Eco1-cohesin interaction, cohesin acetylation, and sister chromatid cohesion establishment.

### Analysing AlphaFold 3 predictions as populations

In the course of our investigations, we realised that AlphaFold 3 predictions do not necessarily settle on a single structure. Rather, repeated runs with multiple independent seeds resulted in populations of predictions that covered a range of structural features. Quantitative analyses of these features resulted in unanticipated biological insight. For example, nicked DNA inside the budding yeast cohesin acetylation complex was found in two configurations, straight, or bent and partially unwound. While the majority of predictions showed bent and partially unwound DNA, alongside an expanded Pds5-DNA interaction surface, the AlphaFold 3 ranking score distribution did not in fact discern the DNA conformations. Relying on just a few model predictions, and choosing the best-ranked model as ground truth, might have led us to miss the striking mode of how the cohesin acetylation complex engages a nicked substrate, which we confirmed by experimental probing of the predicted Pds5-DNA interaction surface.

Another example where a population analysis was instrumental for gaining biological insight was the active state prediction of the cohesin acetylation complex bound to Smc3 and Scc1. In this case, the active state with acetyl-CoA in attack distance from Lys113 was observed in a minority of the predictions. Again, the ranking score did not single out this conformational state. Despite its low frequency, the predicted active state contained distinct structural features, e.g. the Eco1 zinc finger-Scc1 interaction, that our subsequent experiments confirmed. Treating each set of predictions as a distribution proved more informative than selecting a single best model. We expect that other AlphaFold-based investigations of molecular mechanisms can benefit from sampling populations of predictions as a way to explore transient or multi-state assemblies.

### Nicked DNA recognition by the cohesin acetylation complex

Recognition of nicked DNA is not restricted to the cohesin acetylation complex. The replication factor C large subunit Rfc1 similarly recognises nicked DNA by its sharp bend and local unwinding, resolved by cryo-EM^62^. Rfc1 engages both the 3’ and 5’ ends of the nicked strand as well as the exposed single stranded region of the continuous strand, using charge and hydrophobic contacts comparable to those made by Pds5 in the cohesin acetylation complex. While the N-terminal Rfc1 DNA binding site is dispensable for PCNA loading at primer-template junctions^63^, it plays a probable role in DNA structure recognition as part of the DNA damage response to alkylating agents^64^. The PCNA-associated Fen1 flap endonuclease also recognises its substrate by the sharp DNA bend and partial unwinding at both sides of the strand interruption^67^. In this case, binding to the strand interruption directs the Fen1 nuclease active site to the flap overhang.

Other enzymes use alternate strategies for DNA nick recognition. The Poly(ADP-ribose) polymerase PARP-1 detects DNA single-strand breaks and coordinates their repair by transferring ADP-ribose groups from NAD^+^ onto target proteins. In this case, two flexibly connected zinc fingers bind to double stranded regions flanking the nick in a geometry that exploits the extreme deformability of the strand disruption. While nick recognition is achieved differently, the zinc finger arrangement upon nick binding seeds co-operative conformational adaptation of the remaining PARP-1 domains that result in PARP-1 catalytic activation^65^, reminiscent of how conformational adaptations lead to activation of the cohesin acetylation complex. Yet another example for nick recognition is the PCNA-associated DNA Ligase 1, which engages in extensive minor groove interactions that redirect the path of the double helix to expose the nick termini for the strand-joining reaction^66^. The predicted cohesin acetylation complex adds to a number of instances where a DNA strand interruption is recognised to guide downstream molecular events.

#### Coordinating sister chromatid co-entrapment and cohesin acetylation

Okazaki fragment processing intermediates define a window of opportunity for Smc3 acetylation. The spatial proximity of where cohesin co-entraps sister chromatids to these structures might be sufficient to coordinate co-entrapment and acetylation. Circumstantial evidence suggests that co-entrapment and acetylation might be more intimately linked. A number of replisome-associated cohesion establishment factors contribute to efficient sister chromatid cohesion establishment. While their molecular contribution to co-entrapment remains to be understood, they all affect Smc3 acetylation^68^. The cohesin loader complex also contributes to cohesin acetylation in an as yet ununderstood manner^23^. Future experiments will be required to comprehend how these additional factors, directly or indirectly, promote cohesin acetylation. These studies might reveal mechanisms additional to DNA structure dependent Eco1 catalysis, which we have described here.

## Acknowledgements

We thank Daniel Felfoldi and Alessandro Costa for their collaboration, Masashi Minamino for contributing the experiment shown in Figure S6B, Vincent Guacci for the Pds5 and Katsu Shirahige for the ac-Smc3 antibodies, Stephane Mouilleron for discussions and advice, the Fermentation Service in the Crick Structural Biology Science Technology Platform, and all our lab members for discussions and critical reading of the manuscript. This work received support from a Wellcome Trust Investigator Award (220244/Z/20/Z), and from the Francis Crick Institute that receives its core funding from Cancer Research UK, the UKRI Medical Research Council, and the Wellcome Trust (cc2137).

## Author contributions

C.B and F.U. conceived the study. C.B. performed the AlphaFold 3 predictions and their analysis, as well as accompanying experiments, C.B. and F.U. wrote the manuscript.

## Declaration of generative AI and AI-assisted technologies in the manuscript preparation process

Claude Opus 4.8 was used to generate Python code to analyse the AlphaFold 3 models, as described in the Methods. The generated code has been deposited with Zenodo and can be accessed at ‘tbc’.

## Methods

### Yeast strains

Budding yeast *S. cerevisiae* strains used in this study are listed in the Key Resources Table, with genotype details contained in Table S1. Cells were grown in YPD medium at 30 °C it not stated otherwise.

### AlphaFold 3 structure predictions

Nucleoprotein complex structures were predicted using AlphaFold 3^24^ (v3.01 and 3.02) or AlphaFold3_ReD^32^ where specified (red_1.0.0, repulsive bias applied on Pds5 and Eco1 only), on a NVIDIA GPU GH100 80Gb. 5 models each were generated for 75 seeds, if not indicated otherwise. Chain and residue numbers are detailed in Table S2. Note that the position of the nick varies depending on protein composition and complex geometry.

### Contacts and interfaces analyses

Conformationally labile nucleic acid regions (nicked, kinked or melted) were predicted with low pLDDT and high PAE. We therefore implemented distinct contact criteria for protein-protein and protein-DNA interfaces, both gated by AlphaFold 3’s own confidence metrics (interchain PAE and pLDDT). A protein residue pair was deemed in contact if it fulfilled the following criteria: Cβ–Cβ ≤ 8 Å (Cα for glycine) and symmetrised PAE < 15 Å (where PAE is symmetrised as the minimum of the two off-diagonal entries), min(PAE_ij, PAE_ji). A protein-DNA pair was deemed in contact if it fulfilled the following criteria: minimum heavy-atom (non-hydrogen) distance ≤ 4.5 Å to the DNA and Cα pLDDT > 50 on the protein residue. Solvent accessible surface area (SASA) was computed per residue using FreeSASA^69^. For each contact pair, the gated (criteria as above) buried surface area (gated BSA) of a residue was its SASA in isolation minus its SASA when in contact. We report either full chain gated BSA or sub-domain gated BSA, where all residue buried areas between 2 full chains or between sub-domains were summed up. The identities of the sub-domains were: *S. cerevisiae* Eco1 PIP box (14-24), Eco1 zinc finger (30-61), Pds5 patch B (550-580) and Pds5 patch A (1005-1140).

### Bending and melting analysis

For each model, DNA bending was measured from the template-strand phosphate atoms, with the nick position as the vertex. The bend angle is given as 180° minus the internal angle at the vertex, between the first and last phosphate of the strand. DNA melting was reported as the number of template-strand bases lacking a Watson–Crick partner: a template base was scored as unpaired if it did not have at least 2 canonical hydrogen bonds (≤ 3.5 Å) with a base on the other strand.

### Display of representative structural models

We applied a pLDDT > 50 filter on all protein chains shown in our structural models, generated using Pymol. This threshold was lowered to pLDDT > 40 for the *S. pombe* and human complexes, where the ‘YSR’ motif elements showed a pLDDT score between 40 and 50, while nevertheless being confidently placed on Pds5 (PAE < 15).

Protein interfaces were depicted as side chains of one partner and a coloured surface of the other. Mutated residues were highlighted in red.

### Protein-DNA footprints

Gated DNA-protein contacts were computed for the 375 models containing either linear or nicked DNA. Protein and DNA footprints were displayed as the percentage of models showing a specific contact (in case of nicked DNA out of the 229 models that showed bent DNA). The linear DNA assumed different positions amongst the 375 models, so we anchored Eco1 and displayed the relative positions of Pds5 and PCNA.

### Proteins expression and purification

Recombinant Eco1 (wt, PIP box and zinc finger mutants) as well as PCNA and Pds5 were expressed and purified as previously described^23^. Budding yeast cells carrying plasmids expressing either Pds5-ProtA, Smc dimer (Smc1-ProtA+Smc3) or cohesin trimer (Scc1-ProtA+Smc1+Smc3) under the control of *GAL1* promoter were grown in YP+2% raffinose to OD_600_=1. Overexpression was induced by addition of 2% galactose for two hours. Cells were harvested and resuspended in lysis buffer (50 mM HEPES-KOH pH7.5, 300 mM NaCl, 2 mM MgCl_2_, 0.5 mM TCEP, 10% Glycerol, EDTA-free SigmaFast protein inhibitor cocktail), the suspension was dropped into liquid nitrogen, then mechanically broken in a freezer mill under liquid nitrogen. The resulting frozen powder was stored at −70 °C until used.

### ProtA/IgG pulldown

Frozen lysates expressing ProtA-fused proteins were thawed, diluted in lysis buffer and centrifuged at 21,000 x g for 30 minutes at 4 °C. The protein concentration of the clarified supernatant was determined, and 1 - 5 mg of protein were incubated with 30 μl of covalently coupled IgG Dynabeads (NHS Dynabeads + rabbit IgG, coupled according to the manufacturer’s instructions) for 1 hour. The beads were extensively washed with lysis buffer and either purified His6-Eco1 and PCNA, or an *E. coli* lysate containing overexpressed His6-Eco1, were added to the beads and incubated for 30 minutes. The beads were then extensively washed in lysis buffer and the bound proteins eluted in 1.5 x SDS-PAGE sample buffer at 95 °C for 5 minutes. Eluates were separated on 4-12% Tris Glycine Novex gels run in MES/SDS buffer. Proteins were detected either by Coomassie blue staining or western blotting.

### Electrophoretic mobility shift assay

Linear and nicked DNA substrates were prepared by annealing of oligos CB1116 and CB1117 or of oligos CB1116, CB1118 and CB1119 respectively. Oligonucleotide sequences can be found in the Key Resources Table. 0.267 pmol of DNA were incubated for 10 minutes at 25 °C with excess recombinant His6-Eco1 (0 to 40 pmol) in 10 μl of EMSA buffer (25 mM Tris-HCl pH 7.5, 25 mM NaCl, 100 mM potassium glutamate, 0.5 mM TCEP, 50 μg/ml BSA, 5% glycerol). Samples were resolved on Novex 10% TBE gels for 50 minutes at 150 V, stained with SYBR gold and imaged using a BioRad ChemiDoc imaging system.

### Grafix and glycerol gradients

6 ml of 25% glycerol solution (25% glycerol w/v, 50 mM HEPES-KOH pH 7.5, 150 mM NaCl, 0.5 mM TCEP containing crosslinkers (2 mM BS3 and 0.16 mM glutaraldehyde) if indicated, was layered beneath 6 ml of 10% glycerol solution (10% glycerol w/v, 50 mM HEPES pH 7.5, 150 mM NaCl, 0.5 mM TCEP). Gradients were formed using a BioComp gradient station and left at 4 °C until use. 1500 pmol of individual or combined proteins (His6-Eco1, Pds5, PCNA trimers, DNA) were loaded onto the gradients and centrifugated at 4 °C in a Beckman SW41Ti rotor at 38,000 x rpm (183,000 x g) for 19 hours in an Beckman optima ultracentrifuge with acceleration 7 and deceleration 9 settings. The gradient was then harvested from the top. Each 200 μl fraction was quenched with glycine at a final concentration of 100 mM. An aliquot of every second fraction was analysed by SDS-PAGE followed by Coomassie blue staining or western blotting.

### Size exclusion chromatography

600 pmol of Eco1 (wt, PIP box and zf finger mutants) and 600 pmol of PCNA trimers were incubated for 30 minutes at 4 °C, in 60 μl of SEC buffer (50 mM HEPES-KOH pH 7.5, 250 mM potassium glutamate, 1 mM MgCl_2_, 0.5 mM TCEP, 50 mM acetyl-CoA, 3% glycerol), before being separated on a Superdex 200 increase 3.2/300 column, at a flow rate of 20 μl/min using an Äkta pure micro liquid handler. Protein peaks were detected by their UV absorption at 280 nm.

### Analysis of yeast mutants

Cells carrying auxin-inducible degron alleles of *PDS5* (*pds5-iaa7*), *ECO1* (*eco1-iaa7*) and *SCC1* (*scc1-iaa17*) were constructed and complemented with their corresponding either wild type or mutant variants, expressed under control of their endogenous promoters. The ability of the mutant proteins to support cell growth was assessed by spotting 10-fold serial dilutions of a cell suspension at OD_600_=0.5 on YPD plates in the absence or presence of auxin (2 mM indole-3-acetic acid). Plates were incubated at 30 °C and cell growth was assessed after 48 hours. Smc3 acetylation and sister chromatid cohesion were assessed in G2 phase after auxin-mediated depletion of endogenous wild type Pds5, Eco1 or Scc1. Cells were grown in YPD to an OD_600_ = 0.2 and arrested in G1 by addition of 2 μg/mL of α-factor at t = 0, 60 and 120 minutes. After 90 minutes, 2 mM auxin was added to induce protein depletion. Cells were filtered, washed with YP and released in YPD medium containing 2 mM auxin and 24 μg/mL nocodazole. After 90 minutes, cells had reached a mitotic arrest and were harvested to analyse Smc3 acetylation status and sister chromatid cohesion as described previously^70^.

---

**Figure S1.**
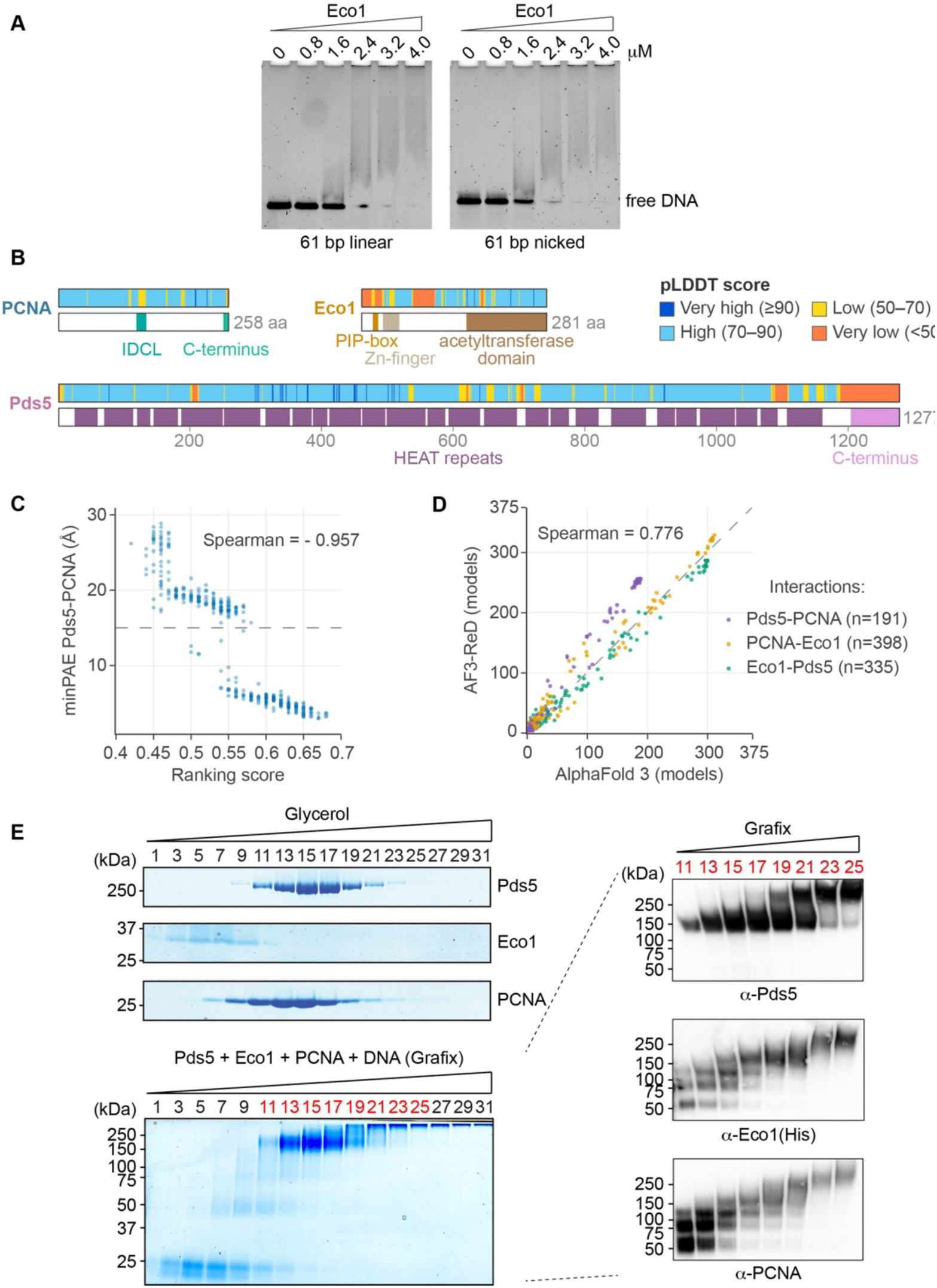
Further characterisation of the predicted and experimental cohesin acetylation complexes. (A) DNA binding by the isolated Eco1 protein. Increasing concentrations of Eco1 were incubated in excess with either a linear or nicked DNA substrate and the resultant Eco1-DNA complexes analysed by native TBE acrylamide gel electrophoresis followed by SYBR Gold staining. (B) Evaluation of AlphaFold 3’s cohesin acetylation complex prediction using the predicted Local Distance Difference Test (pLDDT) score. Linear maps of PCNA, Eco1 and Pds5 are colour coded by their per-residue confidence, next to maps with annotated protein features (IDCL, the PCNA interdomain connecting loop). (C) Correlation plot of the min PAE score at the Pds5-PCNA interface and the AlphaFold 3 ranking score of the 375 analysed models. (D) The number out of 375 how often individual residue pairs between cohesin acetylation complex components (colour-coded) were seen amongst AlphaFold 3 or AF3-ReD predicted cohesin acetylation complexes. Each symbol represents a residue pair. (E) Experimental cohesin acetylation complex assembly. Purified PCNA, Pds5 and Eco1 were either individually subjected to glycerol gradient centrifugation, or mixed in the presence of a nicked DNA template with a one nt gap, and centrifuged through a gradient of the same glycerol concentrations but in the presence of increasing glutaraldehyde and BS3 concentrations (GraFix^33^). Gradients were fractionated and fractions analysed by SDS-PAGE followed by Coomassie Blue staining or Western blotting using the indicated antibodies.

**Figure S2.**
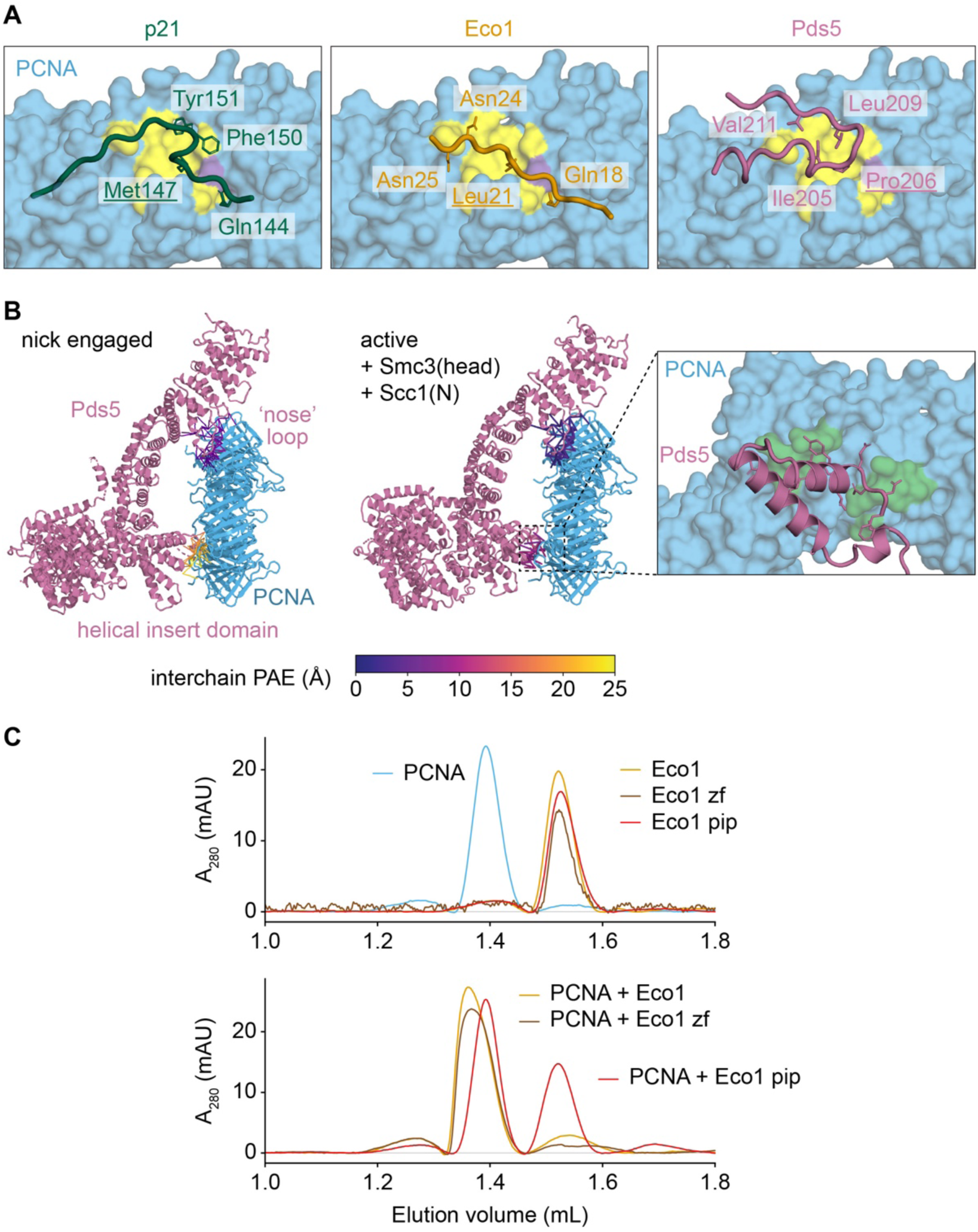
Further characterisation of predicted and experimental protein interfaces. (A) Close up views of the canonical p21 PIP box bound to PCNA^71^, as well as AlphaFold 3 predictions of how the degenerate Eco1 PIP box and the Pds5 ‘nose’ loop interact with the same PCNA surface. The PIP box consensus hydrophobic residues and the residue making an equivalent contact in Pds5 are underlined. (B) Protein interactions at the Pds5 ‘nose’ loop-PCNA and at the Pds5 helical insert domain-PCNA interfaces, colour coded by their PAE scores. Shown are the models from Figure 1A (nick engaged) and Figure 5A (active, + Smc3(head), + Scc1(N). A close-up of the Pds5 helical insert domain interaction with PCNA from Figure 5A is shown alongside. (C) Superdex 200 size-exclusion chromatography analyses of purified PCNA, wild type Eco1, Eco1 with a mutant PIP box (Eco1 pip) and Eco1 with a mutated zinc finger (Eco1 zf). Proteins were analysed individually (top) or after mixing PCNA with one each of the Eco1 variants (bottom).

**Figure S3.**
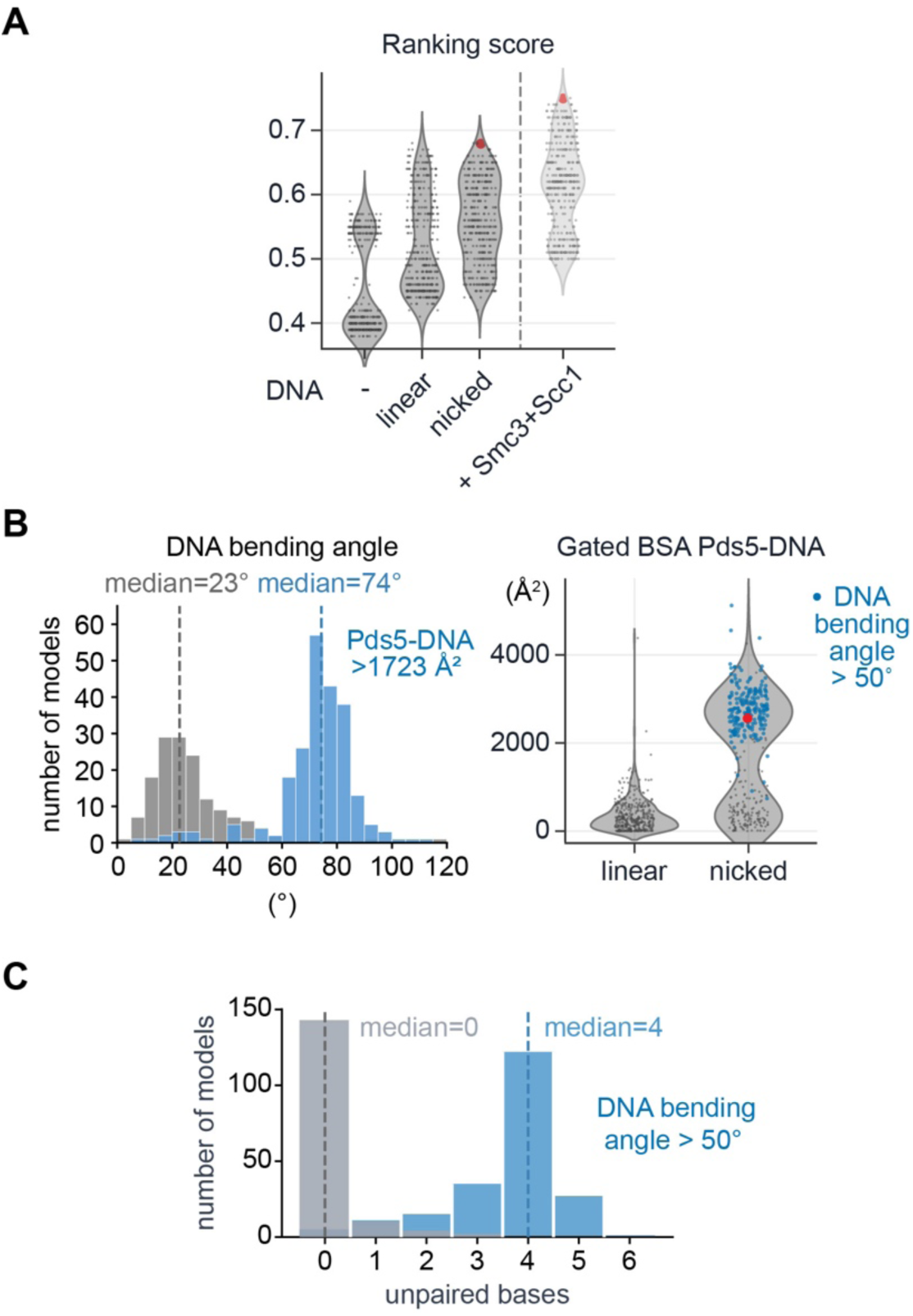
Further characterisation of how the cohesin acetylation complex is predicted to engage with DNA. (A) Violin plots showing the spread of the AlphaFold 3 ranking scores of 375 cohesin acetylation complex models generated in the absence or presence of the indicated DNAs. The ranking scores of 375 cohesin acetylation complex models generated with nicked DNA and in the additional presence of Smc3(head) and Scc1(N) or shown alongside. The models shown in Figures 1A and 5A, respectively, are highlighted red. (B) DNA bending angle distribution amongst 375 cohesin acetylation complex models generated in the presence of nicked DNA, where the blue colour highlights those models with a Pds5-DNA gated buried surface area (BSA) greater than 1723 Å (left). Also shown is the Pds5-DNA BSA spread amongst models generated with linear or nicked DNA, with the blue colour highlighting models with a DNA bending angle of > 50°. (C) Distribution of unpaired base counts at the nick amongst 375 cohesin acetylation complex models generated in the presence of nicked DNA. The blue colour highlights models with a DNA bending angle of > 50°.

**Figure S4.**
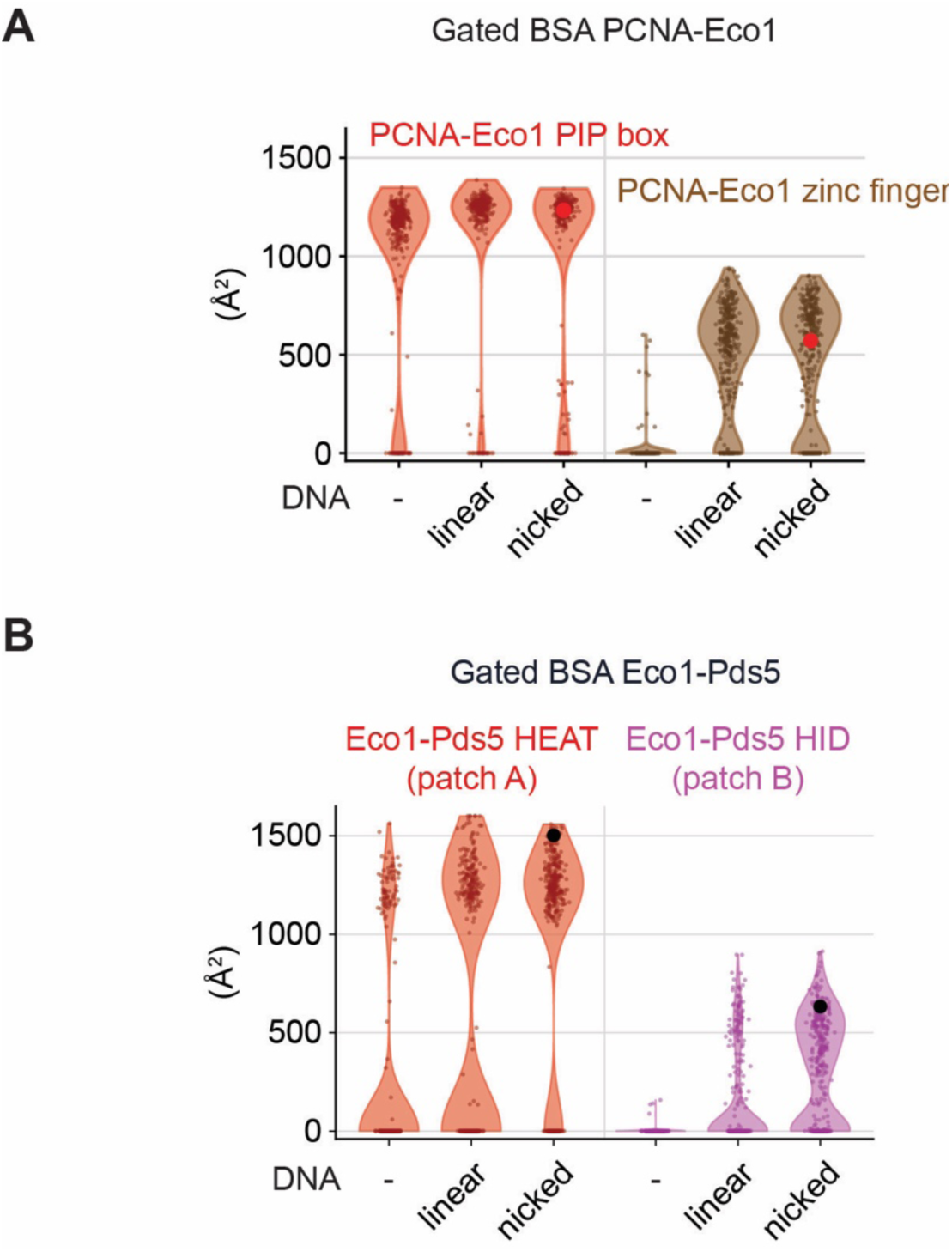
Predictions of how DNA engagement tunes protein interactions within the cohesin acetylation complex. (A) Violin plots of showing the spread of gated buried surface area (BSA) between PCNA and Eco1, separated into PIP box and zinc finger interactions, in 375 cohesin acetylation complex models generate in the absence or presence of the indicated DNAs. The model shown in Figure 1A is highlighted in red. (B) As (**A**) but analysing the Eco1-Pds5 interface, separated into patch A and patch B interactions.

**Figure S5.**
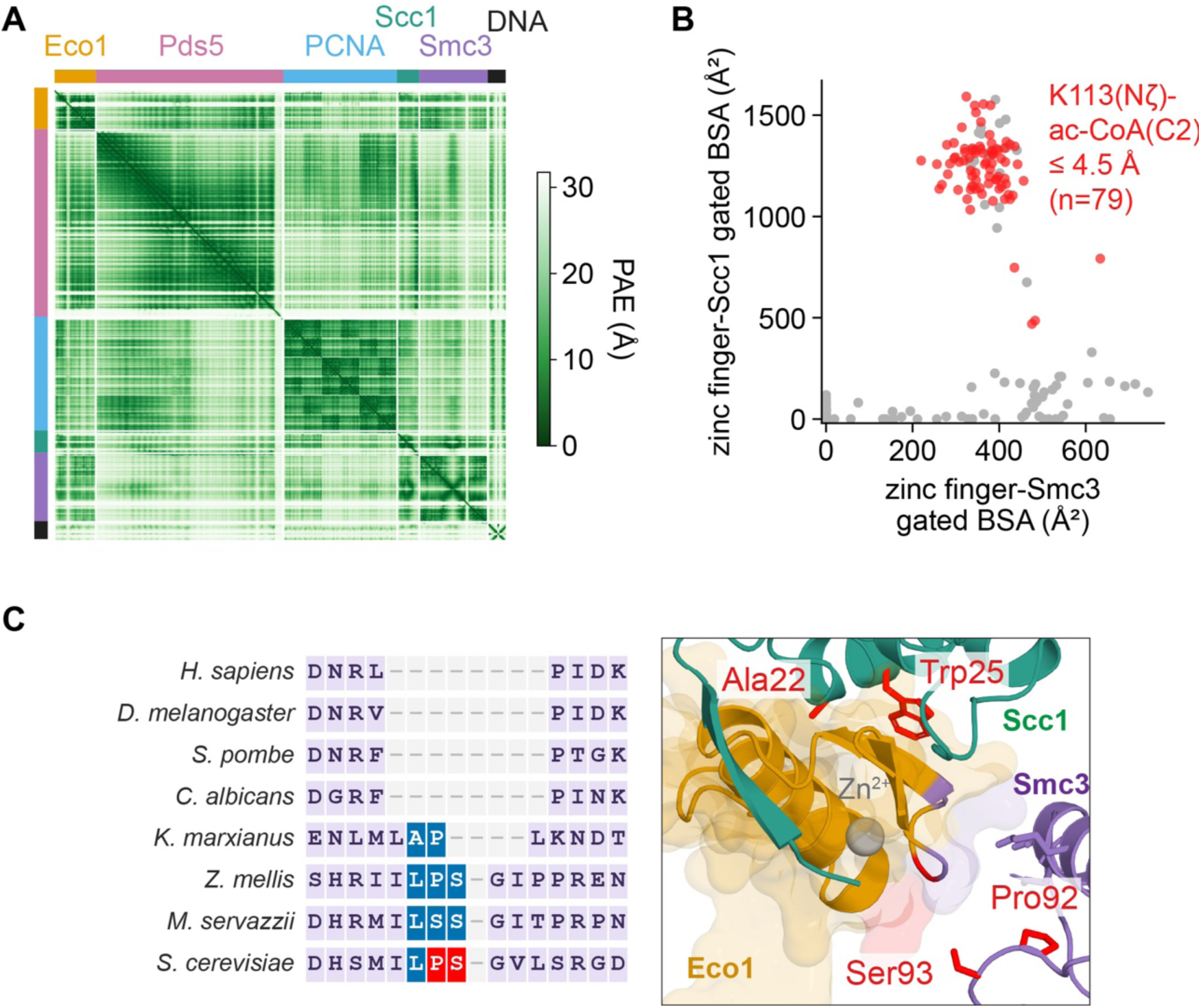
Additional characterisation of the cohesin acetylation complex active state. (A) Predicted aligned error (PAE) matrix of the cohesin acetylation complex model shown in Figure 5A, engaged with Smc3(head) and Scc1(N). (B) The buried surface area between the Eco1 zinc finger and both Smc3 and Scc1 are plotted across the 375 predictions of the cohesin acetylation complex engaged with Smc3(head) and Scc1(N). Colour coded are those models with a distance between the carbonyl carbon of acetyl-CoA and the ζ-nitrogen in the Smc3 acceptor Lys113 < 4.5 Å. (C) Alignment of the amino acid sequence surrounding the Smc3 surface loop containing *S. cerevisiae* Pro92 and Ser93, whose mutation is detrimental for sister chromatid cohesion^45^. The loop is present across budding yeasts but no other eukaryotes. A close-up view of how the surface loop is predicted to engage the Eco1 zinc finger is shown. Scc1 Ala22 and Trp25 are shown for context.

**Figure S6.**
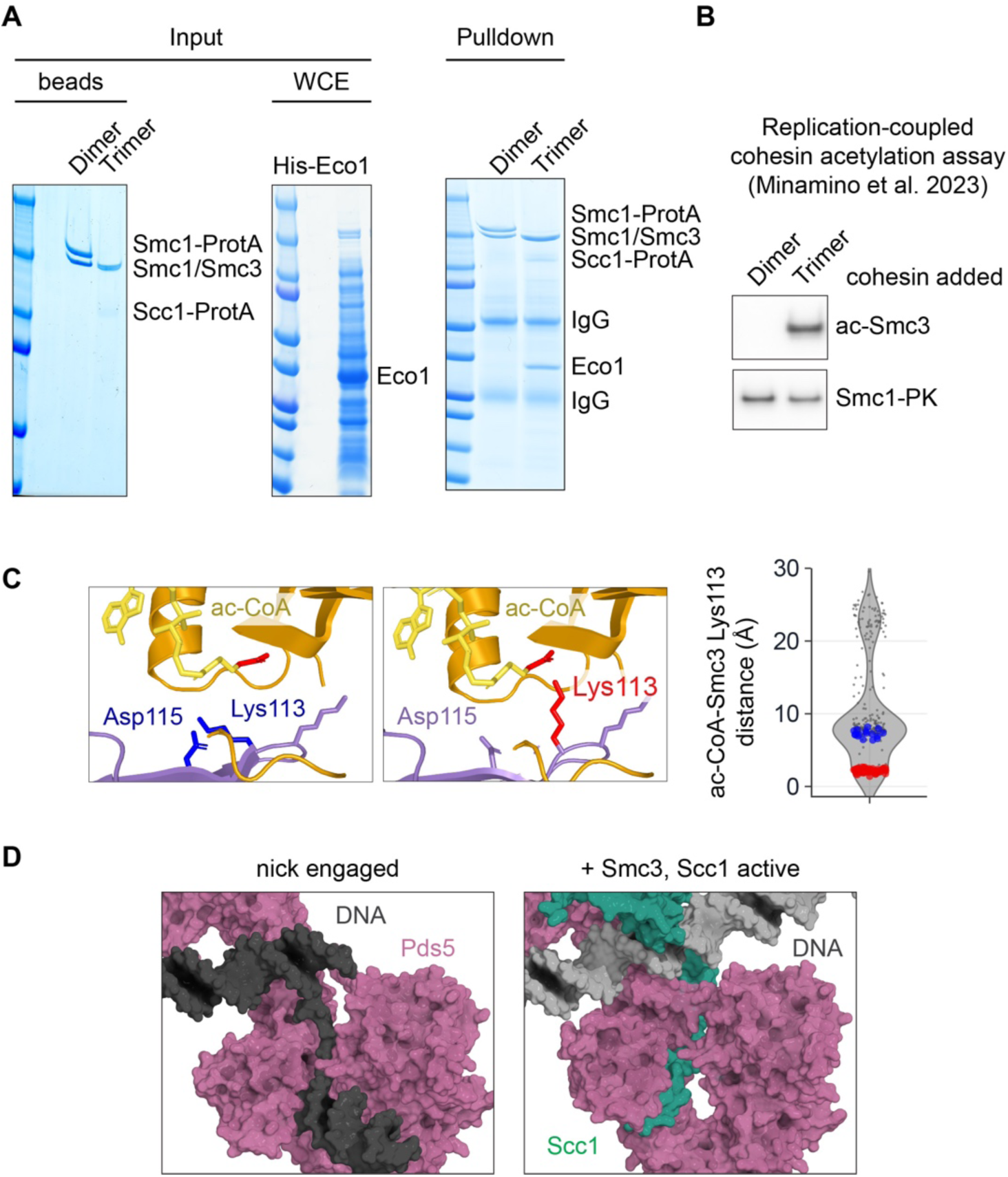
Additional characterisation of the cohesin acetylation complex active state and the role of Scc1 in its assembly. (A) Pull-down of Eco1 by either a Smc1-Smc3 dimer or Smc1-Smc3-Scc1 trimer complex. The complexes, fused to a protein A (ProtA) tag, were immobilised on IgG beads and incubated with bacterial extracts containing overexpressed Eco1. Input and pulldown fractions were resolved by SDS-PAGE, followed by Coomassie Blue staining. (B) A replication-coupled cohesin acetylation assay was performed as described^23^, using DNA polymerase δ for extension of a singly primed template DNA, in the presence of Pds5, and using either the Smc1-Smc3 dimer or Smc1-Smc3-Scc1 trimer as the cohesin substrate. (C) Example close-up views of Smc3 Lys113 and its surrounding in cohesin acetylation complexes capture in their active state, pointing towards the neighbouring Asp115 (blue) or acetyl-CoA (red). The distance between the carbonyl carbon of acetyl-CoA and the ζ-nitrogen in the Smc3 acceptor Lys113 in 375 AlphaFold 3 models is plotted as in Figure 5B, with those models highlighted in which the Eco1 zinc finger engages Scc1 and Smc3, and the orientation of Lys113 towards neighbouring Asp115 (blue) or acetyl-CoA (red) colour coded. (D) Close-up views of Pds5 and DNA in the nick engaged cohesin acetylation complex, compared to a similar view of Pds5, DNA and Scc1 in the cohesin acetylation complex engaged with Smc3(head) and Scc1(N) in its active state.

**Figure S7.**
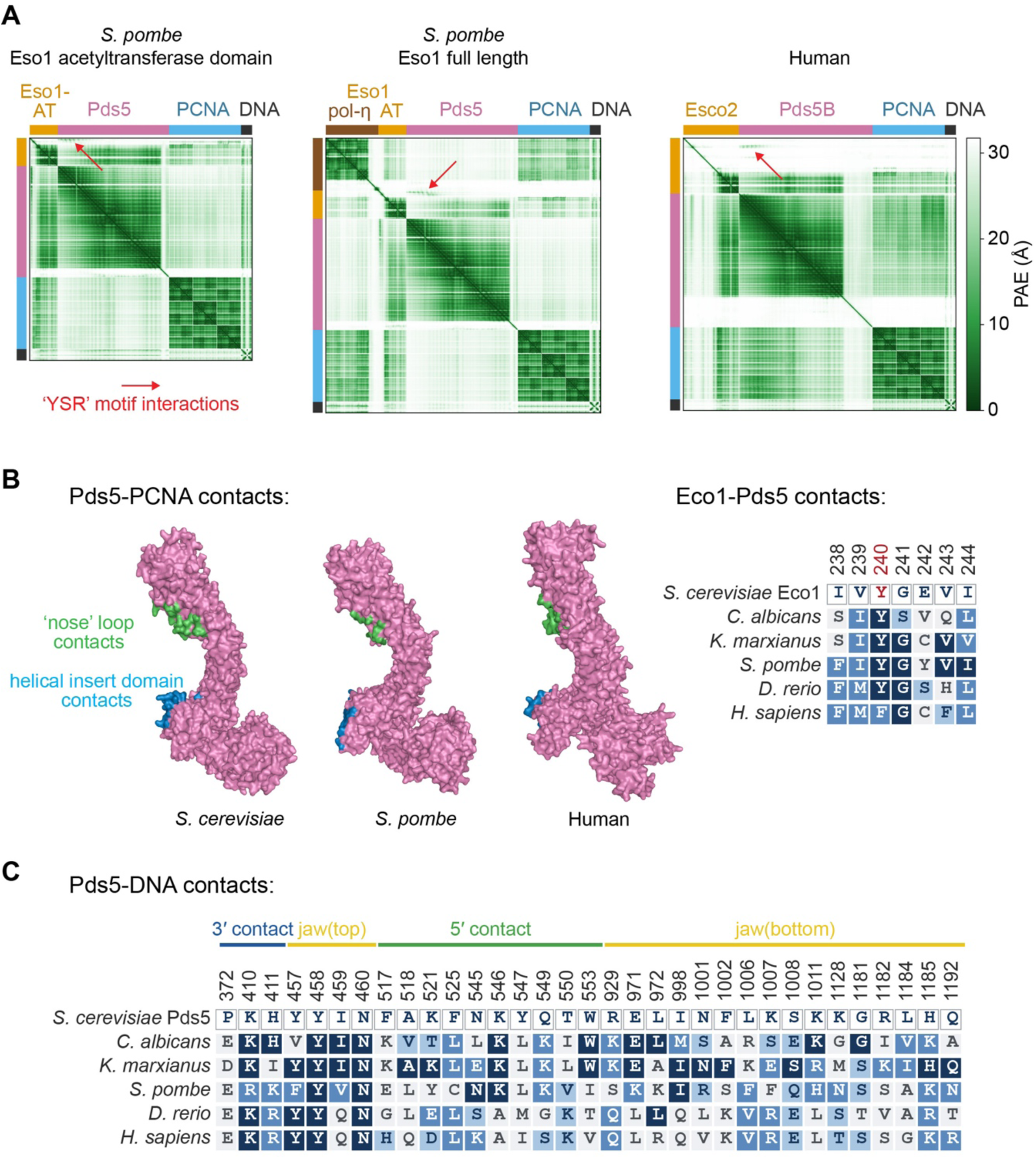
Additional characterisation of the fission yeast and human cohesin acetylation complexes. (A) PAE matrices of the fission yeast and human cohesin acetylation complexes shown in Figures 7A and 7C. (B) Pds5 was extracted from the budding yeast, fission yeast, and human cohesin acetylation complexes shown in Figure 1A and Figures 7A and 7C. Residues contacting PCNA are highlighted. Those in the Pds5 ‘nose’ loop in green, and those in the helical insert domain in blue. Amino acid conservation at the respective acetyltransferase-Pds5 interactions is also depicted. (C) Amino acid conservation of predicted DNA interactions by budding yeast Pds5 is shown, their placements within the 3’ contact, top and bottom parts of the jaw, and 5’ contact are indicated.

